# Systematic mapping of the prokaryotic noncoding sequence space with a genomic language model

**DOI:** 10.64898/2026.10.21.760802

**Authors:** Gwanggyu Sun, Tim Hudelmaier, Joshua C. Gilligan, Gabriel A. Gonzalez, Peter Yin, Fangping Wan, Zijian (Carl) Ma, Matthew G. Durrant, Chiara Ricci-Tam, April Pawluk, Andreas S. Puschnik, Silvana Konermann, Patrick D. Hsu

## Abstract

Prokaryotic genomes encode a vast repertoire of noncoding elements such as ribozymes, riboswitches, and CRISPR RNAs. The sequence-level diversity of these intergenic elements precludes alignment-based clustering that would enable annotation and discovery of novel noncoding RNA families. Here, we show that a genomic language model fine-tuned on prokaryotic intergenic sequences learns representations of noncoding elements from DNA sequence. We used these embeddings to organize 376 million intergenic regions from 1.3 terabase pairs of prokaryotic genomes and metagenomes into 5.14 million clusters. This approach nominated 20,149 clusters of putative structured RNAs with a candidate protein partner. Periodicity in token entropies revealed internal architecture, recovering CRISPR, TIGR and VIPR arrays together with their cognate effector proteins from sequence alone. Targeted clustering of bridge recombinase-adjacent sequences identified bridge RNA variants with longer guide recognition sequences than any characterized ortholog. Finally, we directed teams of AI agents to mine the database under researcher-defined criteria, uncovering divergent riboswitches, ribozymes, and other structured RNAs with biological hypotheses for their function. We release a publicly available database that transforms the dark matter of prokaryotic genomes into an explorable resource for human-agent molecular discovery.

## 1. Introduction

A fundamental goal of the genomics era is to identify and interpret functional biomolecules directly from genome sequence (1, 2). For proteins, this goal is increasingly within reach: structures can now be predicted accurately from sequence (3, 4) and encoding those structures as alignable tokens (5) made it possible to cluster hundreds of millions of predicted structures into millions of families with shared folds and functional annotations (6–8).

However, a large portion of the genomic sequence space remains inaccessible to these approaches. There is currently no structure-aware method to infer evolutionary and functional relationships at scale between divergent noncoding RNAs (ncRNAs) that elude sequence-based alignment (9). Even in prokaryotic genomes, which are compact and densely coding, the noncoding fraction remains far less annotated than the protein-coding regions (10). Prokaryotic ncRNAs, for example, catalyze essential reactions as ribozymes (11), couple metabolite concentrations to gene expression as riboswitches (12), and act as sequence-specific guides for effectors such as nucleases and recombinases (13, 14). Despite their central role in biologically and technologically important processes, the discovery and characterization of ncRNAs remains slow, biased toward model organisms, and largely experimental.

Genomic language models (gLM) such as Evo 2 (15) offer a potential solution. Trained on large and diverse collections of genomes, these models learn representations shaped by evolutionary constraint rather than by annotation. Analogous to how protein language models learn covarying amino acid positions across evolution to infer protein contacts (4), we reasoned that gLM-learned covariation within noncoding sequences could provide the sensitivity necessary to unmask and cluster hidden ncRNAs at scale (**Fig. 1A**). Top-down mapping of the noncoding sequence space would avoid the empirical biases that favor re-discovery or extension of known concepts, while enabling agentic exploration of domain-wide patterns to systematically uncover new ncRNA biology.

**Fig. 1.**
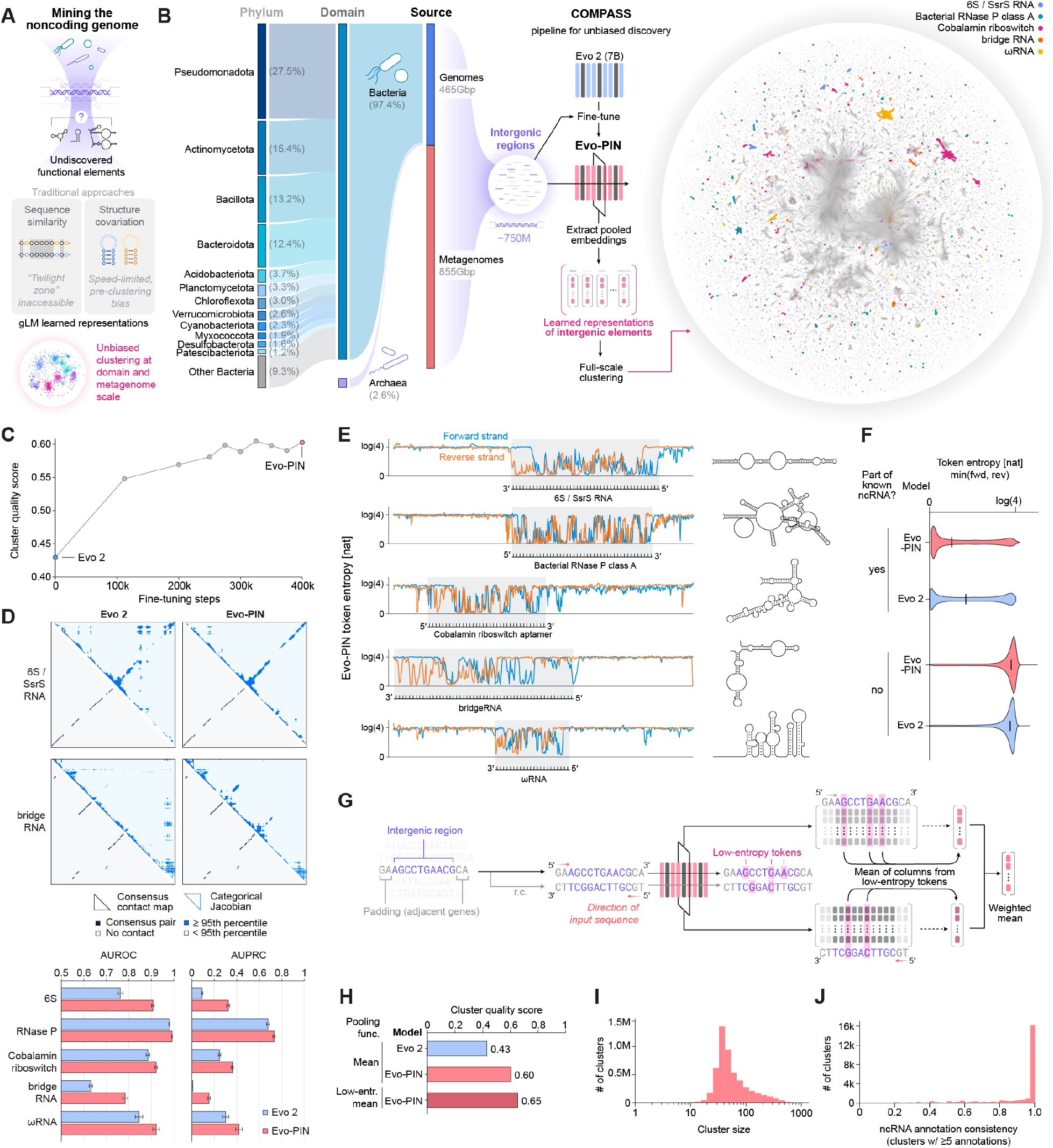
Pooled embeddings from a fine-tuned genomic language model enable unbiased clustering of prokaryotic intergenic regions at scale. (A) Representations of sequences learned by genomic language models can overcome limitations of traditional methods to mine the prokaryotic intergenic sequence space. (B) 750M intergenic regions were parsed from 1.3 Tbp of prokaryotic genomes and metagenomes (left), and used to fine-tune the Evo 2 model. Pooled Evo-PIN embeddings were extracted from intergenic regions, and used to cluster the intergenic regions at scale (center). The resulting clusters of noncoding elements were visualized with UMAP (right), where sequences containing 5 known representative ncRNAs are highlighted. An expanded version with more highlighted ncRNAs is available in Fig. S5. (C) Fine-tuning Evo 2 on prokaryotic intergenic sequences improved the cluster quality score of pooled embeddings, resulting in Evo-PIN at step 400K. The cluster quality score quantifies how well the pooled embeddings from sequences that share the same ncRNA annotation cluster together, with a score of 0 corresponding to shuffled annotations and 1 to perfect clustering. (caption continued on next page) (D) Categorical Jacobians calculated on sequences with known ncRNAs from Evo 2 and Evo-PIN show that fine-tuning improves ncRNA structure predictions. Top: averaged contact scores for 6S/SsrS RNA and bridge RNA. Per-sequence scores were projected onto the consensus columns of each family’s covariance model and averaged across test sequences; upper triangles show consensus column pairs at or above the 95th percentile of averaged score, and lower triangles the consensus base pairs of the known structure. Bottom: area under the precision–recall curve (AUPRC) and the receiver operating characteristic (AUROC) for recovering consensus base pairs from per-sequence contact scores, across five ncRNA families: 6S/SsrS RNA, bacterial RNase P class A, cobalamin riboswitch, bridge RNA, and *ω*RNA. (E) Evo-PIN token entropies along the forward (blue) and reverse (orange) strands of intergenic sequences with known ncRNAs. Token entropies are lower at positions encoding known ncRNAs as indicated with shaded area. Consensus structures of each known ncRNA are shown on the right. (F) Distributions of token entropies for positions inside and outside of known ncRNA annotations, for Evo-PIN and Evo 2. The differences between the two were significantly larger for entropies from Evo-PIN (*p* < 1 × 10^−4^). Black line in violin plots indicates median of token distribution. (G) Schematic of the low-entropy pooling function. Only the embeddings from positions along the sequence with low token entropies (<1.3047) were averaged in this pooling function. (H) Comparison of cluster quality scores for different models and pooling functions as defined in (C). (I) Distribution of cluster sizes (the number of sequences in each cluster) across the 5.14M clusters. (J) Distribution of the consistencies (fraction of the sequences annotated with the most common ncRNA annotation within a cluster) of known ncRNA annotations among the 24,837 clusters with ≥5 sequences uniquely annotated with a known ncRNA.

To this end, we fine-tuned Evo 2 on prokaryotic intergenic sequences, which are otherwise underrepresented in its original training data. We used the embeddings from this model, Evo-PIN (<u>P</u>rokaryotic <u>IN</u>tergenic), to cluster all bacterial and archaeal intergenic sequences top-down in an alignment-and reference-free manner through a pipeline we named COMPASS (<u>COMP</u>rehensive <u>A</u>tlas of <u>S</u>tructured <u>S</u>equences). The resultant 5.14M clusters recapitulate known ncRNA families such as riboswitches, ribozymes, transfer-messenger RNAs, group II introns, CRISPR RNAs and bridge RNAs. Yet, fewer than 1% of the clusters contained known ncRNA annotations, underscoring that the prokaryotic intergenic space has remained underexplored. We demonstrate how feature extraction and clustering from Evo-PIN embeddings, together with enrichment analysis of colocalized proteins, enable discovery and functional predictions across diverse ncRNA families. Periodicity in token entropies uncovered CRISPR, TIGR, and VIPR arrays (16–18), along with uncharacterized repeat arrays. Targeted reclustering of bridge recombinase-adjacent sequences identified bridge RNA variants with longer recognition sequences than any characterized ortholog. Applying this capability across the full COMPASS database requires a throughput that traditional investigation cannot provide. We therefore deployed teams of agents with a “scientist-in-the-loop” framework that read the primary literature, mined public datasets, and performed bioinformatic analyses in parallel, identifying putative divergent riboswitches and hammerhead ribozymes, and uncovering long candidate ncRNAs commonly encoded next to PhoH2 proteins. Finally, we make the COMPASS database openly accessible as an interactive web app (compass.arcinstitute.org) and a Model Context Protocol (MCP) server to accelerate top-down discovery of noncoding biology by the scientific community.

## 2. Results

### 2.1. Clustering of prokaryotic intergenic regions using a fine-tuned Evo 2 model

To build a comprehensive dataset of prokaryotic intergenic regions, we collected 465 Gbp of genomic sequences from 144K bacterial and archaeal species (19) and 855 Gbp of metagenomic sequences (15) (**Fig. 1B, left**). We defined intergenic regions as 10-10,000 bp segments of DNA between annotated protein-coding genes, tRNAs, and rRNAs. After removing redundant sequences, we obtained 750M distinct intergenic sequences with a median length of 113 bp (**Fig. S1A**) and a median GC% of 50.0% (**Fig. S1B**). Each intergenic sequence was searched with covariance models of known prokaryotic ncRNAs from Rfam and other recently discovered ncRNAs (14, 20, 21). Only 0.4% of the sequences had one or more significant hits, and among these only 1.8% had hits for ≥2 distinct ncRNAs (**Fig. S1C**), highlighting both the sparsity of existing ncRNA annotations and the scarcity of regions with two or more distinct elements.

We previously reported Evo 2, a genomic language model trained on DNA sequences from all domains of life (15). To test whether its embeddings group intergenic regions that share a known ncRNA, we defined a cluster quality score on a benchmark subset of intergenic sequences, where 0 corresponds to shuffled annotations and 1 to perfect clustering. Mean-pooled Evo 2 embeddings from these sequences scored 0.43 (**Fig. 1C, S2A**), indicating that the base model already captures ncRNA-relevant features of intergenic sequence.

Because intergenic regions make up only 10% of prokaryotic genomes, and are thus underrepresented in the original Evo 2 pre-training dataset, we fine-tuned the Evo 2 model on the full set of 750M intergenic sequences via continued pre-training (22). This increased the cluster quality score to 0.60, a 39.5% improvement over the base Evo 2 model (**Fig. 1C**). We named this fine-tuned model Evo-PIN (<u>P</u>rokaryotic <u>IN</u>tergenic). To further probe whether Evo-PIN had learned the covarying base patterns within ncRNAs, we computed the categorical Jacobians, which quantify how much a mutation in one position affects the model’s predicted probabilities of bases at a different position (23). Across 5 different ncRNA families, contact maps derived from Evo-PIN’s categorical Jacobians recovered consensus base pairs more accurately than those from Evo 2 (**Fig. 1D, S2B**), indicating that fine-tuning strengthened the model’s representation of the covarying structure.

Since functional ncRNAs often occupy a subsequence within the intergenic region, pooling the embeddings across the full sequence length inevitably dilutes the ncRNA signal. A language model’s token entropy quantifies the level of uncertainty in its prediction for a specific token, with lower entropies signaling more certainty (24). We noticed that Evo-PIN token entropies were lower for positions within a sequence that were annotated with certain ncRNAs (**Fig. 1E**). Globally, Evo-PIN’s median entropies for subsequences encoding known ncRNAs were 73% lower than the entropies outside the ncRNA annotation, while this difference was less pronounced (55%) for Evo 2 (**Fig. 1F**, **S2C**). Additionally, the token entropy of a particular position calculated from the forward strand was often very different from the entropy for the same position calculated from the reverse strand (**Fig. 1E, S2D**), which indicated that the determination of low-entropy positions needed to be strand-specific. From these observations, we devised a new pooling function that selectively averaged the embeddings from positions where the token entropy fell below a threshold value of 1.3047, representing the bottom 1% of entropies from randomized sequences (**Fig. 1G**, **S2E**). Using these low-entropy pooled embeddings resulted in a cluster quality score of 0.65 (8.3% improvement over simple mean-pooled Evo-PIN embeddings) (**Fig. 1H**).

Using an approximate k-nearest-neighbor graph and Leiden clustering (**Fig. S3**, see Methods), we grouped 376M intergenic sequences into 5.14M clusters (**Fig. 1B, right (UMAP); Fig. S4A**). The clusters had a median sequence count of 44, with individual cluster sizes ranging from 1 to 1,363 (**Fig. 1I**). For the 24,837 clusters with ≥5 sequences annotated with known ncRNAs, the identity of the ncRNA annotation was largely consistent within the clusters (0.92 mean consistency) (**Fig. 1J**). To illustrate this, we highlighted multiple known ncRNA families spanning distinct functional classes (**Fig. S5**): catalytic RNAs (bacterial RNase P class A, group II introns), RNAs acting through protein partners (6S/SsrS RNA, tmRNA, SRP RNA), metabolite-sensing riboswitches (TPP, cobalamin, c-di-GMP-I), the tRNA-sensing T-box leader, guide RNAs of mobile elements (bridge RNA, *ω*RNA), and Acido-Lenti-1, an abundant family of unknown function. Of note, each ncRNA family distributes across multiple clusters rather than occupying a single region, reflecting divergence within families and a clustering resolution optimized for the consistency of annotations within each cluster (**Fig. S3G**).

The median pairwise sequence identity between sequences within the same cluster was 28.5% (**Fig. S4B**), well below the “twilight zone” of 50-60% sequence identity where sequence-based alignment algorithms become unreliable for RNAs (9). 24.6% of the clusters had ≥90% of their genomic sequences sourced from the same phylum, while the remainder of clusters were more distributed across multiple phyla (**Fig. S4C**).

We name this pipeline of clustering sequences using gLM embeddings COMPASS. These results demonstrate that COMPASS enables a top-down, alignment- and reference-free organization of the prokaryotic noncoding sequence space.

### 2.2. Cluster-level analysis reveals shared noncoding motifs and colocalized genes

COMPASS groups sequences that share a noncoding element, but does not identify the element itself nor its function. For each cluster, we therefore investigated the conserved RNA structure shared by its sequences and the genes that consistently colocalize with them. Since clustering had used strand-agnostic embeddings, we first reoriented the member sequences of each cluster by iteratively reversing the orientation of each sequence until their forward- and reverse-strand pooled embeddings were mutually consistent (**Fig. 2A, top**), which placed all known ncRNAs within a cluster on a consistent strand with an 80% success rate (**Fig. S6**). We then used CMfinder to identify shared noncoding RNA motifs within each cluster (25), retaining at most one representative motif per cluster after filtering.

**Fig. 2.**
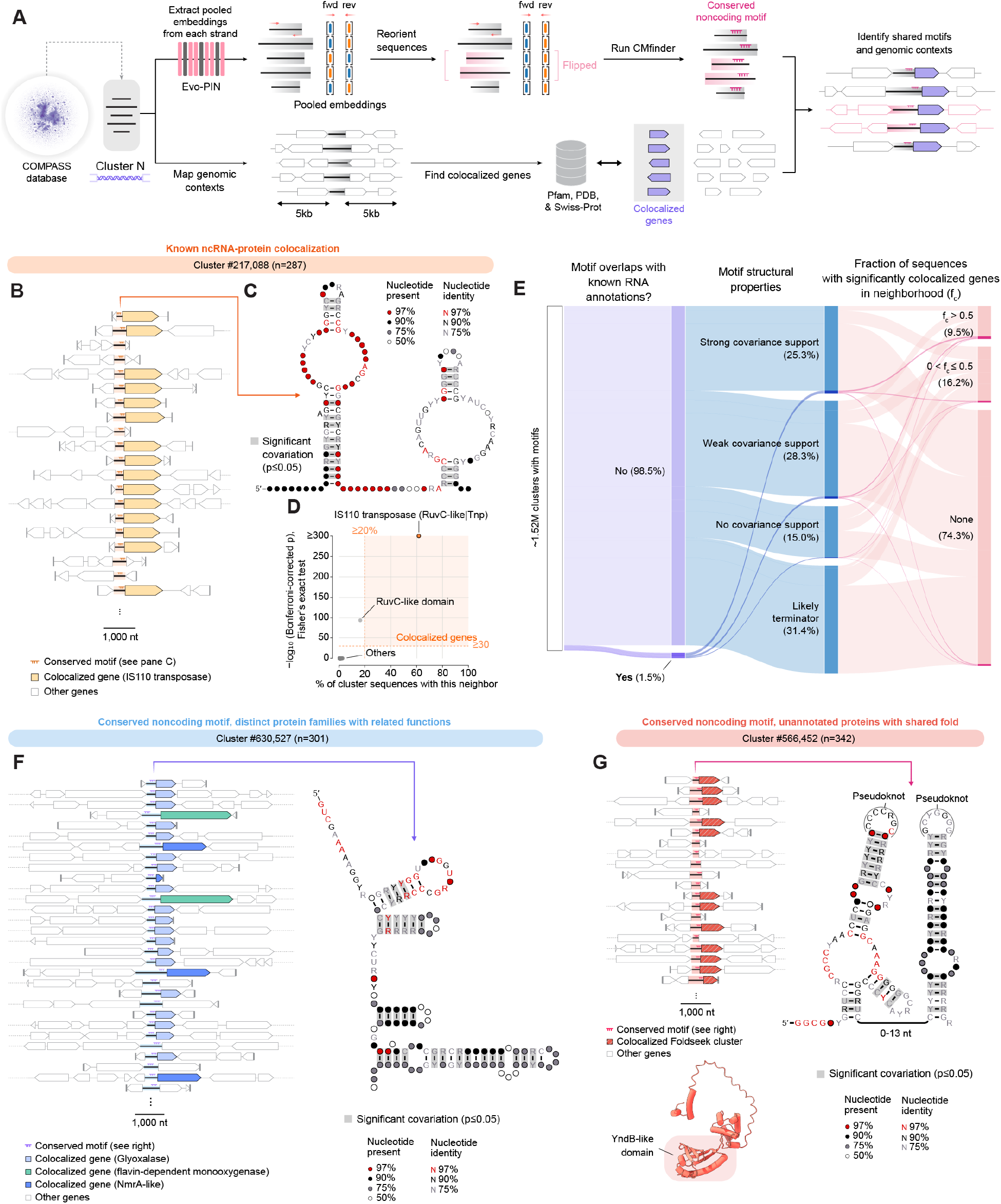
COMPASS clusters share noncoding structures and colocalized genes. (A) Schematics of the work-flow applied to each cluster. Top: Sequences within each cluster are reoriented such that the shared element lies on the same strand in all members. CMfinder is then used to identify a conserved noncoding motif. Bottom: The genomic context of each member sequence is extended by 5 kb in both directions, and the encoded proteins are annotated against Pfam, PDB, and Swiss-Prot to identify genes that colocalize significantly with the cluster. The noncoding motif and its colocalized genes are combined for each cluster. (B) Genomic context of cluster #217,088. Each row is one member locus, aligned on the conserved motif (red); colocalized IS110 transposase genes are shaded orange, all other genes are white. (caption continued on next page) (C) Consensus secondary structure of the cluster #217,088 motif, composed of the two loops characteristic of bridge RNAs. Circle fill denotes the fraction of sequences carrying a nucleotide at that position, letter color denotes nucleotide conservation, and gray shading marks base pairs with significant covariation (*p* 0.05). (D) Enrichment of protein domain architectures in the genomic neighborhood of cluster #217,088. Each point is one architecture, plotted by the percentage of cluster sequences carrying it as a neighbor against the significance of its enrichment (Bonferroni-corrected *p*-value from Fisher’s exact test). Dashed lines mark the thresholds used to call significant colocalization (≥20% of sequences; corrected *p* 10^−30^), and the shaded region indicates architectures meeting both. The IS110 transposase architecture (RuvC-like|Tnp) is the most significantly enriched. (E) Classification of 1.52M CMfinder motif-containing clusters by overlap with known ncRNA annotations, structural properties of the motif and the fraction of cluster sequences with significantly colocalized genes in their neighborhood. (F) Genomic context (left) and motif consensus secondary structure (right) of cluster #630,527 highlighting a conserved noncoding motif adjacent to genes encoding three distinct families of metabolic enzymes. (G) Genomic context (left) and motif consensus secondary structure (right) of cluster #566,452 highlighting a conserved noncoding motif adjacent to genes encoding unannotated proteins with a shared fold. The predicted structure (bottom) of the representative protein from the colocalized Foldseek cluster, with the YndB-like N-terminal domain highlighted.

In prokaryotic genomes, functionally related elements tend to lie close together (2, 26). Genes that consistently colocalize with an uncharacterized noncoding element may hint at its function. We therefore extended the genomic contexts of each sequence by 5 kb in both directions, and annotated any protein-coding genes as well as tRNAs and rRNAs (**Fig. 2A, bottom**).

This combined workflow successfully reconstructed the structures of known ncRNAs and identified their respective protein counterparts. For example, cluster #217,088 is composed of 287 sequences sharing a motif that consistently lies upstream of genes encoding IS110 transposases (**Fig. 2B**). The motif’s secondary structure closely resembled the canonical double-loop structure of bridge RNAs (**Fig. 2C**) (14), and the most significantly enriched protein family in the genetic context of these sequences contained the expected IS110 transposase domain architecture (RuvC-like domain + Tnp domain) (**Fig. 2D**). In a second example, the motif shared across cluster #16,688 displayed key structural features of *ω*RNAs including the nexus pseudoknot, and the sequences were consistently adjacent to a conserved protein fold similar to the OMEGA nickase IsrB, a key partner protein of *ω*RNAs (**Fig. S7A**) (27). These examples demonstrate that our pipeline recovers both structural and contextual hallmarks of these recently discovered ncRNA-guided systems.

Next, we focused on clusters with motifs that do not correspond to any known ncRNA. Of the 1.52M clusters with CMfinder motifs, 1.5% contained motifs that overlap with known ncRNA annotations in at least two sequences and were excluded (**Fig. 2E**). The most common remaining element class was rho-independent transcription terminators, which are characterized by a single GC-rich hairpin followed by a poly-U tract positioned downstream of a protein-coding gene (28). Using their positions relative to their adjacent genes and a structure-based classifier (29), we flagged 476,178 motifs (31.4%) as likely terminators and excluded them from further analysis (**Fig. S7B**). The remaining motifs were tested for covariation to distinguish conserved RNA structures with covarying bases from DNA elements conserved only at the sequence level (30, 31), and classified as having strong, weak or no covariation support (**Fig. 2E**; see Methods). Each cluster was also scored based on the fraction ( *f_c_*) of its sequences that have a significantly colocalized neighboring gene with the constituent noncoding elements (**Fig. 2E**).

In total, 20,149 ncRNA clusters had strong covariation support for their motifs and a clear signature of colocalized, annotated genes ( *f_c_* > 0.5). For example, cluster #630,527 contains a motif immediately upstream of genes that encode glyoxalase, flavin-dependent monooxygenase, or NmrA-like proteins (**Fig. 2F**), suggesting the existence of a common 5^′^ leader sequence shared by these enzymes. Many other motifs were colocalized with proteins of unknown function, such as the motif for cluster #378,112, positioned directly upstream of genes containing DUF736, a largely uncharacterized domain (**Fig. S7C**). For such clusters where the colocalized protein functions are known, it supports the generation of more specific hypotheses about ncRNA function. In cases where the protein is uncharacterized, the ncRNA features may provide information on the molecular function of the protein, such as formation of a ribonucleoprotein complex or RNA-based regulation of its expression (32).

Other clusters with strong covariation support had no annotated colocalized genes. For some, applying Foldseek (5) to the encoded proteins within a ±5 kb window revealed a shared protein fold that was not annotated by Pfam. In cluster #566,452, for example, this fold was located directly downstream of the candidate ncRNA motif (**Fig. 2G**). The structure of the N-terminal domain of the protein was similar to the structure of the uncharacterized protein YndB from *B. subtilis*, which is presumed to function in stress response (33). Other clusters did not colocalize significantly with any genes. For example, sequences in cluster #580,639 shared an extensive motif with 57 predicted base pairs in its structure, yet neither annotated protein-coding genes nor shared folds colocalized significantly (**Fig. S7D**).

Together, these analyses assign to each cluster a predicted secondary structure, a set of significantly colocalized genes, and quantitative support for each. Importantly, these associations were obtained without reference to existing annotation or prior selection of candidate regions.

### 2.3. Periodic token entropy patterns reveal clusters of CRISPR, TIGR, and VIPR arrays

Many known RNA-guided systems are encoded as tandem arrays in which a conserved repeat alternates with a variable segment. In CRISPR arrays, for example, conserved direct repeats are interleaved by variable spacers that confer sequence specificity on their cognate Cas proteins (17). We reasoned that this architecture should produce a periodic signature in Evo-PIN token entropies, with low entropy across the conserved repeats and high entropy across variable sequences. We confirmed that the Evo-PIN token entropies along CRISPR arrays display periodic patterns that match known annotations (**Fig. S8A**) and that sequences containing known CRISPR direct repeats cluster together in our pipeline (**Fig. S8B**). Given these observations, we investigated whether conserved periodic entropy patterns within a cluster can be used to mine for repeat arrays more systematically.

To do this, we autocorrelated each token entropy track against itself at every lag *p*, averaged the resulting profiles across each cluster, and took local maxima of the averaged profile at nonzero *p* as evidence of a repeating pattern of period *p* (**Fig. 3A**). This identified 1,672 clusters (365K intergenic sequences) with a consistent period across their members. The dominant period length was sharply trimodal, with 96.1% of clusters exhibiting periods of 60-79 nt, 18 nt, or 5 nt, matching the spacings of known CRISPR, TIGR, and VIPR arrays, respectively (**Fig. 3B**).

**Fig. 3.**
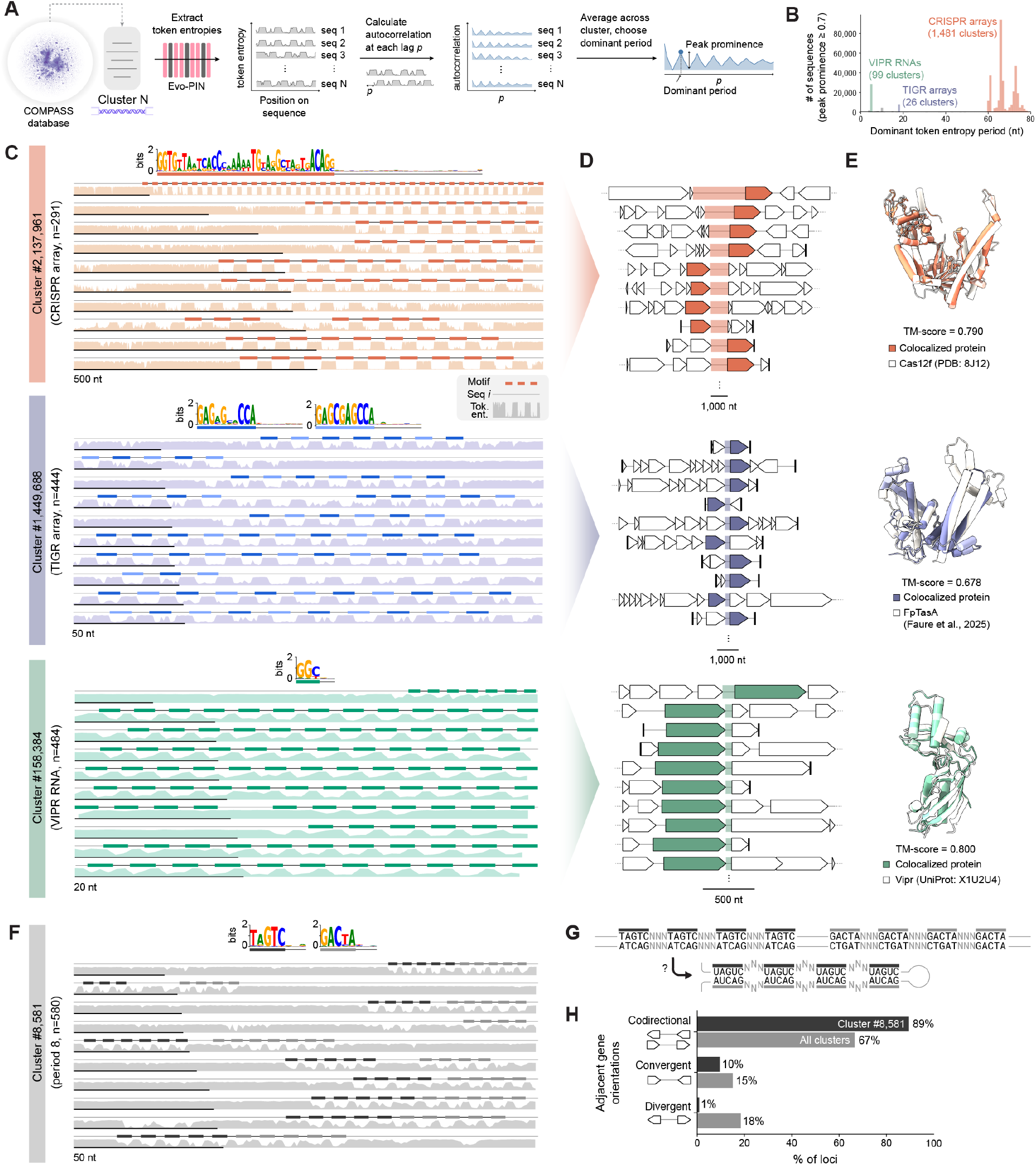
Periodic token entropy patterns identify tandem repeat arrays associated with RNA-guided systems. (A) Schematic of the periodicity analysis. Evo-PIN token entropies are extracted for every member sequence of a cluster, the autocorrelation of each entropy track is computed at every lag *p*, and the resulting profiles are averaged across the cluster. The dominant period is the lag of the most prominent local maximum of the averaged profile, and the topographic prominence of that peak is the cluster’s periodicity score. (B) Distribution of dominant token entropy periods across the 1,672 clusters with a periodicity score ≥ 0.7. Colors denote the bands corresponding to VIPR RNAs (5 nt), TIGR arrays (18 nt), and CRISPR arrays (60-79 nt). (C) Representative clusters from each band: #2,137,961 (CRISPR array), #1,449,688 (TIGR array), and #158,384 (VIPR RNA). Each row is one member sequence, showing its extent (black line), its token entropy track (shaded), and the positions of detected repeat motifs (bars). The sequence logo of the consensus repeat is shown for each repeat array family. (caption continued on next page) (D) Genomic contexts of member loci from the same three clusters. Each row is one locus, with the intergenic region containing the array shaded, the enriched colocalized protein colored, and all other genes white. (E) Superposition of the ESMFold-predicted structure of the most abundant colocalized protein fold from each cluster (colored) onto the known effector of the corresponding system (white): Cas12f (PDB 8J12), FpTasA (18), and Vipr (UniProt X1U2U4). Query-normed TM-scores are indicated. (F) Cluster #8,581 (n = 580), with a dominant period of 8 nt, displayed as in (C). The two logos show the fixed pentamers of the two arrays. (G) Repeat architecture of cluster #8,581, in which an array of fixed TAGTC pentamers separated by variable trimers is followed by a reverse-complemented copy of the same array. Prediction of the “beads-on-a-string” secondary structure that such an element would adopt if transcribed. (H) Comparison of the relative orientation of the two genes flanking each locus for cluster #8,581 with all clusters.

As expected, CRISPR array signatures were the most common of the three, with 1,481 clusters comprising 317K intergenic sequences, compared with 99 VIPR-like and 26 TIGR-like clusters. Cluster #2,137,961 exemplifies the long direct repeats and spacers of a CRISPR array, whereas TIGR arrays (e.g. cluster #1,449,688) display shorter alternating direct repeats (18), and VIPR arrays (e.g. cluster #158,384) have an alternating pattern of GGY/NN motifs (16) (**Fig. 3C**). For all three, the most abundant protein fold in the flanking genomic context matched the system’s known effector: Cas12f, FpTasA, and Vipr, respectively (**Fig. 3D, 3E**).

Further analysis of the CRISPR array-like clusters (34) showed that the sum of the lengths of the direct repeat and the spacer for the same array (which should match the entropy period *p*) formed a trimodal distribution centered around 61, 66, and 73 nt (**Fig. 3B**, **S9A**), driven mostly by uneven distributions in the direct repeat lengths. The lengths of the spacer and repeat, as well as the sequences of the repeats, were more similar for sequences within the same cluster than for random pairs of sequences (**Fig. S9B**). For each sequence, we predicted the CRISPR-Cas subtype for each locus using the identities of the proteins around the array (35). Most clusters had internally consistent type and subtype calls (**Fig. S9C**) despite the sequences being clustered using only embeddings from the intergenic sequences, confirming that the arrays are largely coupled to the identity of the CRISPR-Cas system (36).

In the Vipr-vrRNA complex, two adjacent Vipr protomers lock each repeating unit of the GGY trinucleotides in place, exposing the variable NN dinucleotides to pair with the target. The same alternating signature of fixed trinucleotides and variable dinucleotides was recently used to characterize VIPR arrays from Evo 2 entropy patterns (16), independently corroborating the entropy-based signal we exploit here. A closer examination of the 99 clusters with entropy periodicity at *p* = 5 revealed that most clusters are composed of sequences with the canonical GGY/NN motifs (16), but 21 clusters spanning 6,318 sequences carried dominant 5-mer motifs that do not match this pattern (**Fig. S10A**). These motifs included alternative fixed trinucleotide patterns (e.g. GAC/NN or CAC/NN) as well as motifs degenerate at one position within the trinucleotide (e.g. WGT/NN or TGH/NN), as in the canonical GGY. In all 21 clusters, the non-canonical motifs repeated ≥10 times, and a distinctive shared protein fold was in close proximity to the sequences (**Fig. S10B**), with structural similarity to described Vipr proteins (**Fig. S10C**). This strongly suggests that they represent divergent VIPR systems with non-canonical repeat units.

Finally, only 66 (3.9%) of the periodic entropy clusters fell outside the three periodicity bands of 60-79 nt, 18 nt, and 5 nt. The largest group, 32 clusters at period 10, contained runs of 4-6 adenines or thymines spaced every 10-11 nt, consistent with A-tracts associated with intrinsic DNA curvature (37) rather than structured RNA elements (**Fig. S11A**). Among the remaining clusters were #8,581 and #156,623, with dominant periods of 8 and 7, respectively (**Fig. 3F, S11B**). In both, an array of fixed n-mers alternating with variable 3-mers was followed by a reverse-complemented copy of the same array, with only the fixed n-mer repeats reverse-complementary between them. If transcribed, such an element would be expected to fold into a beads-on-a-string structure, with variable loops separated by a fixed stem (**Fig. 3G**). Neither cluster colocalized significantly with any annotated protein or conserved fold, but the orientations of the two neighboring genes to the loci were predominantly codirectional for both clusters (**Fig. 3H, S11C**). Their genomic sequences were mostly derived from phyla Chloroflexota (#8,581) and Patescibacteriota (#156,623), respectively. Together, these results establish periodic token entropy as a signature of tandem repeat architecture, sufficient to recover known RNA-guided families and to nominate array-type candidates without an identified candidate effector.

### 2.4. Targeted clustering of bridge RNA loci reveals novel orthologs with longer guide pairings

So far, we have used Evo-PIN representations to survey the intergenic sequence space globally, identifying shared noncoding elements through their conserved structures, colocalized genes, and periodic sequence patterns. We next asked whether the same representations could resolve variation within a single ncRNA family based on a specific molecular characteristic. For this purpose, we chose to investigate bridge RNAs (bRNA), the bispecific guide RNAs for bridge recombinase systems. In our previous work, we have identified many variants of bRNAs with diverse secondary structures, with some orthologs having putative longer guide sequences than the canonical IS621 bRNA (14, 38). We sought to discover these diverse orthologs systematically with an embedding-based approach that does not require extensive manual alignment of sequences flanking bridge recombinase proteins.

To specifically yield clusters of bRNA variants, we subsampled intergenic sequences that are adjacent to bridge recombinases, and clustered them using their Evo-PIN embeddings. From each cluster, we built a structural model for the candidate bRNA, and analyzed the potential target and donor guides within the RNAs. The sum of the lengths of these guides is the total programmable nucleotides (TPN) of each bRNA, a proxy for the specificity that the RNA could provide as a genome engineering tool (**Fig. 4A, Methods**). In total, we predicted the TPN for 32,190 bRNAs using 86 different covariance models created from the clusters of the sequence subset (**Fig. 4B**). The predicted secondary structures of many covariance models visibly deviate from the RNA structure described for the canonical IS621 bRNA, yet show clear covariation with their target and donor DNA sequences, validating the ability of Evo-PIN embeddings to organize diverse sets of noncoding sequences within a single family (**Fig. S12A-F**).

**Fig. 4.**
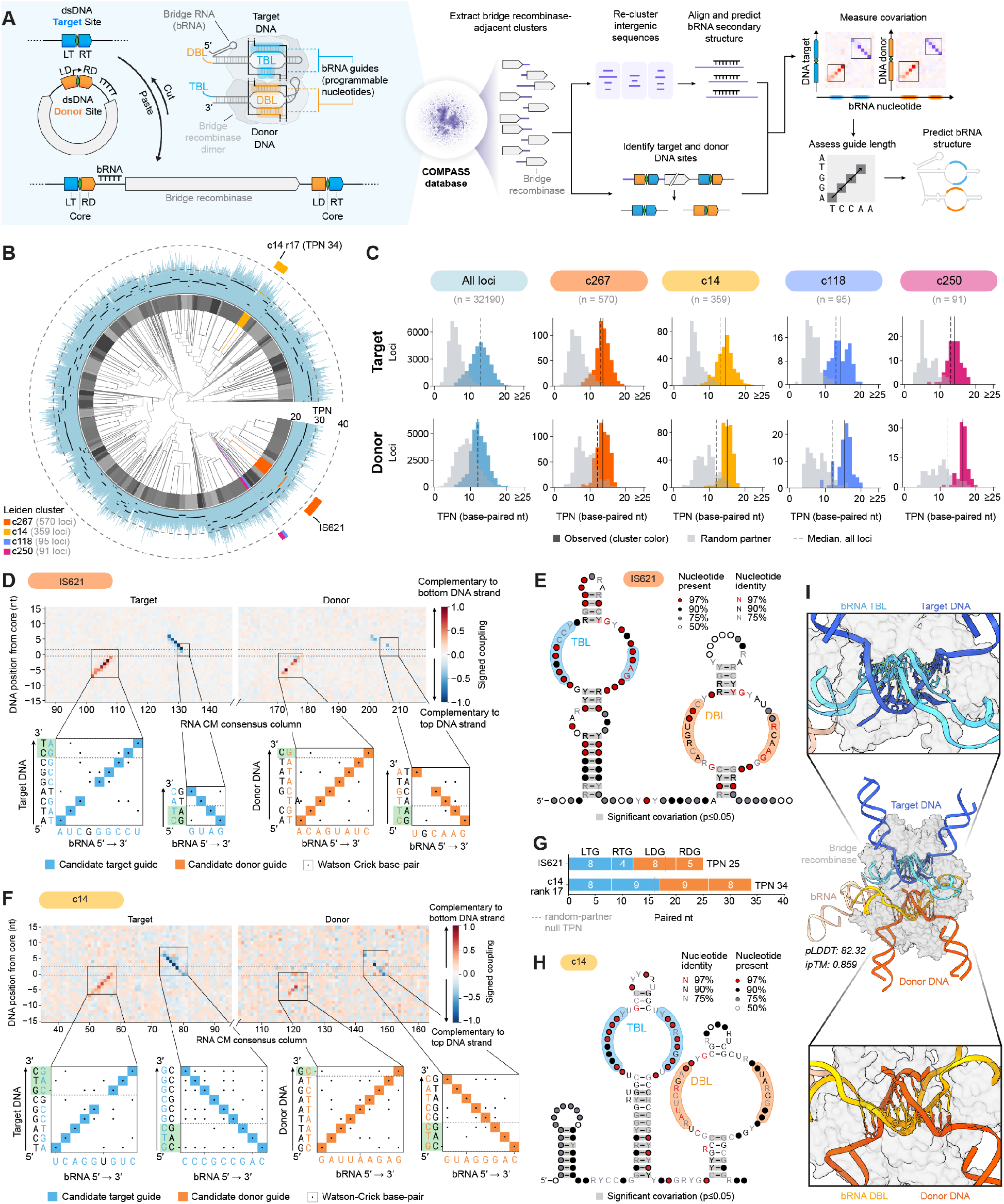
Targeted clustering of bridge recombinase loci identifies variants with longer bridge RNA recognition sequences. (A) Left: Bridge recombinases catalyze recombination between a target and a donor DNA site, each specified by base pairing with a dedicated loop of the bridge RNA (bRNA), the target-binding loop (TBL) and donor-binding loop (DBL), such that recombination joins the left target (LT) and right donor (RD) sequences around a shared core. Right: Workflow for bridge recombinase mining. Clusters enriched for adjacent bridge recombinase genes were extracted from the full database and re-clustered; a covariance model was built and refined for each resulting cluster; element boundaries were used to reconstruct target and donor sites; and covariation between bRNA and DNA positions followed by local alignments was used to call guide <u>lengths. (caption continued on next page)</u>(B) Clustering of bRNAs based on Evo-PIN embeddings. The grayscale ring denotes individual Leiden clusters. Outer blue bars show the total programmable nucleotides (TPN) predicted for each locus with the black line indicating the median TPN for a given cluster. Three representative clusters with high median TPN as well as the prototypical IS621-containing cluster are highlighted in colors. (C) Distribution of predicted TPNs for target(top) and donor-(bottom) binding loops across all loci for the four selected clusters. Colored bars show observed values; gray bars show the null obtained by scoring each bRNA against a randomly assigned target and donor pair of a different bRNA matching the same covariance model. Dashed lines mark the median across all loci. Solid lines mark the median in a specific cluster. (D) Covariation between bRNA and DNA positions for cluster 267. Top: signed coupling between each RNA consensus column and each DNA position relative to the core, for the target and donor site; positive and negative values indicate complementarity to the bottom and top DNA strand, respectively. Boxes mark the four called guide regions. Bottom: predicted base pairing of IS621 within each boxed region, with Watson–Crick pairs indicated. (E) Consensus secondary structure of the cluster 267 bRNA, with the predicted IS621 target- and donor-binding loops shaded. Circle fill indicates the fraction of sequences with a nucleotide at that position, and letter color indicates nucleotide conservation. (F) As in (D), for cluster 14. (G) Predicted cumulative paired nucleotides across the four guide segments for IS621 and a cluster 14 member. (H) As in (E), for cluster 14. (I) Structural model of a cluster 14 bRNA, bridge recombinase, and target and donor DNA, predicted by cofolding. Insets show the target-binding loop paired with target DNA (top) and the donor-binding loop paired with donor DNA (bottom).

Across the 32,190 loci, the predicted TPN ranged from 12 to 40, with a median of 25. The previously characterized IS621 bRNA from *Escherichia coli* was found in cluster 267 along with all other bRNAs encoded next to IS621 recombinases. Cluster 267 had a median TPN of 27 (**Fig. 4C**), close to the experimentally determined TPN of 28 for the IS621 bRNA (14). Using the bRNA model of cluster 267 to predict the guide regions of the *E. coli* IS621 bRNA recapitulated the previously validated base pairing with the exception of 2 base pairs in the DBL (**Fig. 4D, 4E**). Encouragingly for our search for high-TPN bRNAs, 6,113 (19.0%) of the 32,190 analyzed RNAs were predicted to have a TPN greater than 28 (**Fig. 4C**).

Among the 6 clusters with the highest median TPN of 30, cluster 14 covered the most intergenic loci (359), and 68% of its bRNAs were predicted to have a higher TPN than IS621. The predicted secondary structure for this cluster mirrored the predicted secondary structure of the IS621 bRNA (**Fig. 4E, 4H**). A representative locus within cluster 14 contained a bRNA with a predicted TPN of 34 (17 base pairs for both the target and donor; 95th percentile TPN within cluster 14) (**Fig. 4F, 4G**). Cofolding the structures of the predicted bRNA with the associated recombinase protein, target DNA, and donor DNA yielded a structure where all 34 predicted base pairings between the guides and the target and donor DNAs were visible (**Fig. 4I**) (39).

In sum, this illustrates that the same Evo-PIN embeddings used to survey the entire noncoding space can be focused on a single family to resolve variation within it, both recovering the experimentally validated guide segments of the IS621 bRNA as well as nominating previously uncharacterized bRNA orthologs with compelling molecular characteristics.

### 2.5. Discovery of novel candidate noncoding RNAs through researcher–agent collaboration

COMPASS resolves the prokaryotic intergenic sequence space into 5.14M clusters, the vast majority of which lack any ncRNA annotations. While covariance models identify members of established RNA families, divergent sequences can escape detection despite retaining functional motifs, structural features, or conserved genomic associations. Recovering these candidates requires integrating biological evidence beyond the sequence and structural profiles captured by the covariance models. We investigated whether scientific agents could scale this process of biological discovery, autonomously developing computational analyses and synthe-sizing evidence across modalities under researcher guidance. Using a custom agentic harness, we developed a collaborative researcher–agent discovery workflow supporting parallel hypothesis generation, multimodal investigation, and independent review (**Fig. 5A**).

**Fig. 5.**
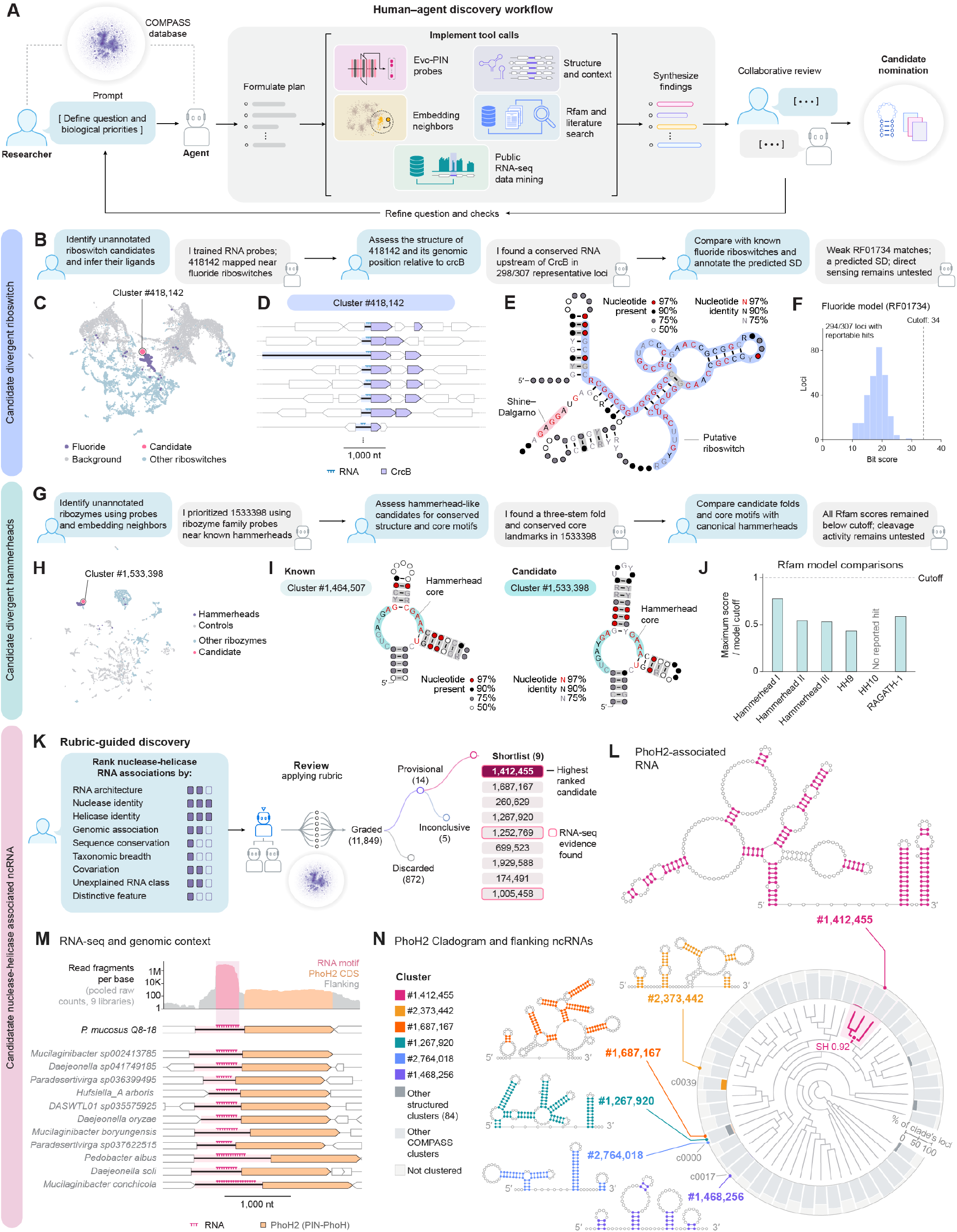
Agentic mining of the COMPASS database nominates novel noncoding RNA candidates. (A) Schematic of the researcher–agent discovery workflow. A researcher defines biological objectives, while an agent formulates investigative plans, executes parallel tool calls, synthesizes evidence, and iteratively refines hypotheses through collaborative review. (B) Representative researcher–agent interactions illustrating the discovery and subsequent investigation of a candidate divergent fluoride riboswitch. (C) Projection of Evo-PIN embeddings showing the position of candidate cluster #418,142 relative to known fluoride riboswitches, other riboswitch families, and background sequences. (D) Genomic neighborhoods showing the conserved colocalization of sequences of cluster #418,142 with co-oriented *crcB* fluoride transporter genes. (E) Predicted secondary structure of the candidate fluoride riboswitch, highlighting conserved nucleotides, base-pairing patterns, and a putative Shine–Dalgarno sequence. (F) Distribution of Rfam fluoride riboswitch covariance model (RF01734) scores across 294 loci of cluster #418,142 with reportable hits. Scores remain below the annotation threshold (dashed line). (G) Representative researcher–agent interactions illustrating the identification and structural investigation of a divergent hammerhead ribozyme candidate. (H) Projection of Evo-PIN embeddings highlighting the position of candidate cluster #1,533,398 relative to annotated hammerhead ribozymes, other ribozyme families, and background sequences. (I) Comparison of predicted secondary structures for a known hammerhead ribozyme (cluster #1,464,507) and candidate cluster #1,533,398, highlighting the conserved three-stem architecture and catalytic core. (J) Maximum Rfam covariance model scores for cluster #1,533,398, normalized by the corresponding family-specific annotation thresholds. All six hammerhead models remain below their respective cutoffs. (K) Rubric-guided discovery of previously uncharacterized nuclease–helicase-associated ncRNAs. A lead agent coordinates parallel investigations integrating RNA architecture, protein identity, genomic association, sequence conservation, and structural evidence. Independent agent review of 11,849 cluster–motif hypotheses yields 14 provisional candidates, including nine shortlisted candidates and the highest-ranked cluster #1,412,455. (L) Predicted secondary structure of the PhoH2-associated RNA candidate #1,412,455, highlighting conserved base-pairing interactions supported by 75% covariance. (M) Genomic organization of candidate PhoH2-associated ncRNAs across representative bacterial loci and RNA-seq coverage from nine pooled *Pedobacter mucosus* Q8-18 libraries, showing transcriptional coverage across the candidate RNA. (N) Phylogenetic distribution of PhoH2 homologs and their associated structured RNAs. Colored branches and representative secondary structures highlight distinct RNA architectures associated with different PhoH2 lineages, including the Bacteroidota-enriched subtree containing cluster #1,412,455.

We first deployed this workflow to identify novel riboswitch candidates. In conversation with the researcher, the agent developed, trained, and deployed embedding-based linear probes across 29 ligand classes (**Fig. 5B**) (40). After scoring all unannotated clusters, the agent prioritized cluster #418,142 as a candidate fluoride riboswitch based on high probe scores (**Fig. 5C, S13A-B**). Additional evidence was derived from a con-served genomic colocalization with the CrcB fluoride ion channel, as well as a predicted consensus secondary structure containing a putative riboswitch aptamer (**Fig. 5D-E**). This cluster represents a potentially divergent family of fluoride riboswitches as none of its 307 members met the score cutoff for the canonical Rfam model (RF01734) (**Fig. 5F**). Using *in silico* saturation mutagenesis (ISM) to systematically substitute each nucleotide and measure the changes in probe scores (41, 42), the agents revealed that the probe is sensitive to changes in the conserved core motifs of the novel riboswitch candidate, despite the regions flanking the cores diverging from the known family (**Fig. S13C-E**). To investigate whether this capability extended beyond ligand-binding RNAs, we applied the same approach to catalytic ribozymes (**Fig. 5G**). The agent developed and trained probes across ten ribozyme classes, consistently outperforming a nearest-neighbor baseline in distinguishing ribozymes from GC-matched and flanking-region negatives (**Fig. S13F**). It identified cluster #1,533,398 as a candidate hammerhead ribozyme and recovered a three-stem RNA architecture containing the conserved 5^′^-CUGA-3^′^ catalytic motif (**Fig. 5H-I, S13G-H**). By contrast, none of the six existing hammerhead Rfam models reached their annotation thresholds across its 152 loci (**Fig. 5J**). Together, these examples demonstrate the agent’s ability to construct analytical tools and pursue complementary biological evidence to recover candidates beyond the recognition limits of existing family models.

The previous tasks were focused and interactive. We next tested whether investigation could be scaled across thousands of hypotheses and pursued autonomously by agents after the researcher defined an evidence rubric to standardize scoring across agent-generated hypotheses. We applied this framework to investigate previously uncharacterized structured RNAs associated with proteins containing nuclease and helicase domains. A lead agent orchestrated parallel subagents to investigate 14,722 clusters, evaluating 11,849 distinct cluster–motif hypotheses using evidence spanning RNA architecture, genomic context, protein identity, and evolutionary conservation (**Fig. 5K**). Hierarchical evaluations and independent agent reviews advanced 297 hypotheses to detailed investigation, yielding 14 provisional candidates, of which cluster #1,412,455 alone satisfied the complete researcher-defined scoring rubric.

Cluster #1,412,455 carries a conserved multistem RNA structure that lacks any known-family annotation (**Fig. 5L**). Its genomic context revealed a recurrent association with PhoH2, a protein with PIN nuclease and PhoH helicase domains. Mycobacterial PhoH2 homologs have previously been shown to couple RNA helicase and ribonuclease activities (43). PhoH2-encoding genes are located adjacent to 148 of 326 loci in the cluster and within 5 kb of 228 loci (**Fig. 5M**). The agent further analyzed nine publicly available *Pedobacter mucosus* RNA-seq datasets, finding concentrated transcriptional coverage across the candidate motif (**Fig. 5M**). Phylogenetic analysis uncovered distinct, structurally conserved RNA architectures in other clusters that are associated with other PhoH2 lineages (**Fig. 5N**), suggesting lineage-specific RNA partners across the divergent PhoH2 family (**Fig. S14**).

While the molecular functions of these ncRNAs require experimental validation, we demonstrate that the presented researcher–agent collaborations assembled a traceable body of evidence and identified concrete directions for experimental testing. Collectively, these results mark a shift from computational annotation towards scalable scientific investigation, in which agents develop their own analyses, interrogate thousands of hypotheses, and synthesize heterogeneous evidence to uncover ncRNA candidates inaccessible to conventional family-model searches alone.

## 3. Discussion

A scalable, global map of the biological sequence space serves as a foundation to systematically interrogate the “dark matter” in sequence data that is accumulating far faster than it can be interpreted. In this work, we demonstrate that Evo-PIN, a fine-tuned genomic language model trained only on annotation-free DNA sequences, can be used to build a global map of the prokaryotic intergenic sequence space that is not limited by any alignment requirements or scale, through a pipeline we name COMPASS. Earlier *de novo* surveys of this noncoding space uncovered thousands of candidate structured ncRNAs (44–49), but each required selecting a subset of sequences for examination: either by phylogeny, by sequence similarity, or by adjacency to a particular class of protein. Surveying the full intergenic space without pre-selection removes these constraints, and enables systematic ncRNA discovery across a broad set of contexts.

From Evo-PIN, we extracted four distinct, biologically relevant readouts that each contributed to the characterization of these sequences across contexts. Pooled embeddings grouped intergenic regions that share a particular type of noncoding element. Token entropies marked the position of that element within a locus, enabling us to selectively pool the representations from informative positions. Periodicity in the token entropies exposed internal architecture, recovering CRISPR, TIGR and VIPR arrays in a reference-free manner. Categorical Jacobians recovered consensus base pairs, indicating that the representation reflects evolutionary constraint on structure rather than conservation of sequence. These four readouts were obtained from a single model without task-specific supervision, highlighting generalist capabilities of the model that extend beyond the pre-training objective. Further readouts likely remain accessible from the same model, and systematically mining gLM internals will likely constitute a general route for even richer biological discovery.

A few key steps still separate an organized sequence space from the ability to assign molecular function. Clustering requires no annotation, but our interpretation of the clusters is biased towards signatures that are already recognizable, such as covariation within secondary structures and periodic repeat arrays. Colocalization nominates likely protein partners without establishing how the noncoding elements engage with them. Finally, the ncRNA candidates reported here are computational predictions, and experimental testing remains the rate-limiting step in any genome mining campaign. Our database provides a path to prioritize those experiments. Each cluster carries a predicted secondary structure with covariation statistics, a set of significantly colocalized genes, a taxonomic distribution and, where sufficient data exist, transcriptional evidence. Experimental effort can therefore be directed at the candidates for which the assembled evidence is strongest, and designed around a property shared across a family, a set of variants differing in one predicted parameter, or an RNA and its nominated partner in combination.

The natural extension of this work would be to apply the same approach to the noncoding regions of viral and eukaryotic genomes. Viral genomes are known to harbor many structured ncRNAs, including internal ribosome entry sites (IRES), exonuclease-resistant RNA (xrRNA) motifs, and ribozymes (50, 51). The compactness of these genomes, and the resulting overlaps between genomic elements may make it more challenging to reliably parse relevant noncoding segments for fine-tuning and clustering. Eukaryotic ncRNA annotations remain concentrated in a small number of animal, plant, and fungal model systems, leaving the ncRNA repertoires of other eukaryotic lineages largely unexplored (52). Several features of large eukaryotic genomes would nonetheless require the approach to be adapted. Intergenic regions are orders of magnitude longer, such that pooling the embeddings from a full intergenic segment as a single unit may not be a feasible strategy in these genomes. The relative lack of sequence diversity in eukaryotic genomes may also constrain the statistical power of covariation-based structure validation (30). Finally, the relative lack of conserved gene neighborhoods in most eukaryotes would limit the genomic context signal we relied on to generate functional hypotheses in prokaryotes.

A database of this scale requires modes of exploration that scale with it. Only 1.5% of motif-bearing COMPASS clusters correspond to an existing ncRNA annotation, emphasizing the extent of unknowns in the prokaryotic noncoding sequence space. We demonstrated two approaches in which agents conducted highly parallelized investigations of the database under researcher direction: probe-based search over the embedding space, which deorphanized divergent members of characterized ncRNA classes that fall below the thresholds of their own covariance models; and rubric-guided batch investigation, which nominated structured RNAs with no family assignment by assembling evidence from genomic context, sequence conservation, and public transcriptomic data. We anticipate that agents will guide larger portions of candidate selection and hypothesis generation as AI model capabilities grow, eventually reaching autonomous levels. To facilitate this, we make the COMPASS database openly available to the scientific community through an interactive web application for direct exploration and an MCP server for agentic uses. Together, COMPASS provides a global map of noncoding sequences and agent-driven interrogation offers a route to connecting noncoding sequence diversity to molecular function at a scale that was previously inaccessible.

## 4. Methods

### 4.1. Extracting sequences of intergenic regions from prokaryotic genomes and metagenomes

We acquired 465 Gbp of genomic sequences of 143,614 representative prokaryotic species (136,646 bacteria and 6,968 archaea) from the Genome Taxonomy Database (GTDB) (19) release 226 (April 2025). The protein-coding gene boundaries from the same genomes, called by Prodigal v2.6.3 (53) and provided as part of the GTDB release, were used to first extract the sequences of inter-CDS regions from each genome. Metagenomic sequences of 855 Gbp in total length were acquired from OpenGenome2 (15), and the protein-coding gene boundaries were likewise called by running Prodigal v2.6.3. Any inter-CDS sequences shorter than 10 nt or longer than 10,000 nt were discarded at this stage. Then, we used tRNAscan-SE 2.0.12 (54) and Barrnap 0.9 (55) to annotate tRNAs and rRNAs, respectively, within each inter-CDS sequence. For both tools, we selected options to use bacterial RNA models to call the RNAs in sequences that were parsed from bacterial genomes and metagenomes, while archaeal RNA model options were used to call RNAs originating from archaeal genomes. We used the default score cutoffs for both tools. If tRNAs and/or rRNAs were predicted within an inter-CDS region, the sequence was further segmented into multiple loci based on the boundaries of the called tRNAs and/or rRNAs. Any resulting segments shorter than 10 nt were discarded. Each of these split segments, or any inter-CDS locus without predicted tRNAs or rRNAs, was defined as an intergenic region in this work. Any segments bounded by contig ends were discarded for genomic data but retained for metagenomic data. This process resulted in a total of 1.01B intergenic regions (192.4 Gbp) from both sources, with 305M intergenic loci (45.9 Gbp) coming from genomic sequences and 708M intergenic loci (146.5 Gbp) from metagenomic sequences.

During this extraction step, we also prepared a “padded” set of intergenic sequences where each intergenic region was extended by 20 nt on either side along the source contig into the neighboring genes to serve as padding sequences for embedding extraction and token entropy calculations. We also logged the source genomes of each intergenic sequence, the relative locations of each CDS, tRNA, and rRNA gene located within 5 kb of either end of the intergenic sequence, and the amino acid sequences encoded by the protein-coding genes within the context.

To remove homologous sets of intergenic sequences, we used MMseqs2 version 18.8cc5c (56, 57) to cluster the 1.01B unpadded intergenic segments based on sequence homology. The linclust option of the tool was used to form clusters of sequences with 70% sequence identity at 80% coverage. This resulted in 750M sequence clusters in total, and the longest sequence from each cluster was selected as the representative sequence.

### 4.2. Annotation of known noncoding RNAs in intergenic regions

To annotate known noncoding RNAs (ncRNAs) within the intergenic regions, we searched each of the representative intergenic sequences against a library of covariance models (CMs) using Infernal’s cmsearch (version 1.1.5, (58)). The library comprised mainly of CMs from Rfam 15.0 (20), with CMs for *ω*RNAs (21) and bridgeRNAs (14) added from their respective sources. cmsearch was run with the effective search-space size (-Z) set to 152426.435Mb, the summed length of the 750M representative intergenic sequences. Raw hits were then filtered and consolidated as follows.

Hits were first restricted to those with a score greater than or equal to the model’s gathering threshold (GA), or, if the GA is not given for the model, an e-value 1 10^−3^. For the “strict” set of annotations, any truncated hits or hits to miRNA CMs or CMs whose Rfam seed alignments were composed entirely of eukaryotic sequences (annotated as “Eukaryota (100.0%)” in the Rfam taxonomy table (https://github.com/Rfam/rfam-taxonomy)) were excluded. For this set, we also used a greedy, score-prioritized procedure to resolve any overlapping hits – any overlapping hits were sorted by bitscore in descending order, and each hit was retained only if its coordinate interval did not overlap any previously retained (higher-scoring) hit on that target. This yielded a non-overlapping set of highest-confidence annotations per locus. For the “permissive” set of annotations, no additional filters were applied except for the GA and e-value thresholds. The strict set was used to evaluate the performance of the embedding-based clustering method. The permissive set was used to evaluate whether the motifs of a given cluster overlap with the annotations of a known ncRNA. We do note that many tRNA and rRNA annotations remain in these regions, despite the tRNA and rRNA annotations being parsed out from the original sequences. Many of these hits are fragmented hits, or divergent tRNAs or rRNAs that did not get filtered out by the default score filters of the tools.

### 4.3. Calculation of mean pooled Evo 2 or Evo-PIN embeddings from intergenic regions

Padded intergenic sequences were passed through Evo 2 or Evo-PIN, and the models’ embeddings were extracted from the pre-normalization layer of block 25, which yielded a matrix of dimension *L* 40, *D*, where *L* is the length of the intergenic sequence and *D* is the embedding dimension (4,096 for both models) (**Fig. S2A**). Since in autoregressive language models the embedding at position *i* reflects the model’s internal representation used to predict the nucleotide at position *i* + 1, we reassigned the embedding at position *i* to position *i* + 1, thereby aligning each embedding with the nucleotide it is used to predict. We then masked out the embeddings aligned with the padding regions on both sides and averaged the remaining embeddings along the length of the sequence, which yielded a vector of size D. We repeated the same procedure for the reverse complement of the padded sequence to calculate the pooled embedding for the reverse strand. The final pooled embedding (vector of size D) was an average of these two strand-specific embeddings.

### 4.4. Calculation of cluster quality scores

To calculate cluster quality scores on a benchmark subset, we randomly sampled 10M sequences from the full set of 750M representative sequences, along with their strict annotation table for known ncRNAs. We then extracted pooled embeddings from each sequence using Evo 2 7B or Evo-PIN. For each of the 466 ncRNA families with at least 3 annotations among the 10M subsampled sequences, we identified the set *S_l_* = *i*: *y_i_* = *l* of sequences that are uniquely annotated with the ncRNA family *l*. For each of the sequences with the ncRNA, we examined its pooled embedding *l_i_*, and identified its *k* = 16 nearest neighbors in the pooled embedding space (*N_i_ k*) measured by cosine similarity. From this neighborhood graph, we calculated a normalized precision score for each ncRNA family as follows:

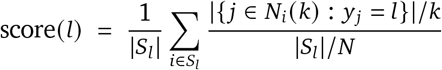

Here, *N* is the total number of uniquely annotated sequences in the 10M benchmark subset. This score searches the neighborhood of a sequence in the pooled embedding space, and calculates the fraction of neighbors that share the same ncRNA annotation as the given sequence, normalized by what is expected by random chance given the distributions of the annotations. Given this score formulation, a purely random neighborhood graph has an expected score of 1, and any scores higher than 1 would indicate that sequences with the same annotations are enriched in each other’s neighborhoods in the pooled embedding space. After calculating the scores for each ncRNA family, we calculated an average over the set of ncRNA families for the aggregate score:

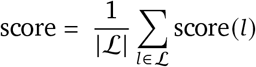

To make the score easier to interpret, we divided the values by the maximum score that can be achieved on the dataset. We calculated the maximum score by assigning each family of ncRNAs only neighbors of their own family. For families with less than k neighbors, we distribute the remaining edges randomly across other families. The normalized score is calculated as follows (**Fig. 1C**, **1H**, **S2F**):

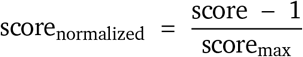

In this way, the score ranges from a minimum of 0, equivalent to a random neighborhood graph, to 1, which is equivalent to perfect nearest neighbor assignment.

### 4.5. Fine-tuning of Evo 2 7B with intergenic sequences

The fine-tuning dataset was constructed by taking all unpadded, representative intergenic sequences (750M) and their reverse complements. After randomizing the order of the sequences, all parameters of the Evo 2 7B (1M context model) were fine-tuned on this dataset using a causal language modeling objective with a context length of 2,048 tokens and naive sequence packing, with training examples being separated by an <EOS> token. Training was distributed across 32 H100 GPUs using data parallelism with ZeRO stage 1 (59), with a global batch size of 524,288 tokens. We used AdamW (60) (*β*_1_ = 0.9, *β*_2_ = 0.95, *ε* = 1*e* 8, weight decay 0.1) with gradient clipping at a global ℓ2 norm of 1.0. The learning rate had a linear warm-up over 57.5K steps (5%) to a peak learning rate of 5 × 10^−5^, followed by cosine decay to 5 × 10^−6^. Intermediate checkpoints were scored using the mean pooled embeddings and the cluster quality scores described above. Fine-tuning was stopped after 400K steps, at which the model has been trained on 35% of the full training dataset, after observing convergence in the cluster quality score. The weights at 400K steps are referred to as Evo-PIN.

### 4.6. Calculation of categorical Jacobians for sequences with known ncRNA annotations

We evaluated categorical Jacobians (23) on five ncRNA families: 6S/SsrS RNA, bacterial RNase P class A, the cobalamin riboswitch, bridge RNA, and *ω*RNA. For each family, we collected all representative intergenic sequences with a strict annotation for that family and discarded sequences with hits on both strands. Each remaining sequence was reverse-complemented where necessary so that the ncRNA was on the forward strand, and was cropped to the ncRNA hit plus up to 100 nt of flanking sequence on each side; for sequences with multiple hits, the highest-scoring hit was used. To obtain a non-redundant evaluation set, we clustered the cropped sequences of each family with MMseqs2 easy-cluster (90% sequence identity, 80% coverage) and randomly sampled 300 sequences with a single hit, each from a different cluster. All categorical Jacobian results were computed on these 300 sequences per family. To obtain ground-truth base pairs, we aligned the hit region of each sequence to the CM used for its annotation with Infernal’s cmalign and projected the consensus base pairs onto the sequence, keeping a pair only if both of its columns were aligned to nucleotides. Pairs with *j − i* < 4 were excluded. Fifty-two *ω*RNA sequences had no projected base pair, leaving 248 *ω*RNA sequences for evaluation.

We calculated categorical Jacobians for Evo 2 7B and Evo-PIN. For a sequence of length *L*, each position *i* was substituted with each of the three alternative nucleotides, and the resulting change in the unnormalized A/C/G/T logits at every position *j* relative to the unmodified sequence was recorded as a tensor *J* of size*L* 4 *L* 4. Because both models are autoregressive, a substitution at position *i* only affects predictions at downstream positions. We therefore ran each substituted sequence on both the forward strand, which provided the entries with *j* > *i*, and its reverse complement, which provided the entries with *j* < *i* after mapping positions and nucleotides back to the forward strand. A single nucleotide was prepended to each input so that the first position was also predicted. *J* was mean-centered along each of its four axes and symmetrized by averaging the (*i*, *a*, *j*, *b*) and ( *j*, *b*, *i*, *a*) entries. The contact score of each pair (*i*, *j*) was the Frobenius norm of its 4 × 4 block; after setting the diagonal to zero, we applied the average product correction (APC) and then set entries with |*i* − *j*| < 2 to zero.

Classification metrics were computed on the aligned region of each sequence, i.e., the annotated ncRNA hit interval (the segment aligned by cmalign), excluding the flanking sequence. Positives were the projected consensus base pairs, and negatives were all other pairs *i*, *j* with *j i* 4 and both nucleotides in the aligned region. We calculated the AUPRC and AUROC of the contact scores for each sequence, and reported the mean over sequences with 95% percentile-bootstrap confidence intervals from 2,000 resamples of sequences (**Fig. 1D**).

For the contact maps in **Fig. 1D** and **S2B**, the contact scores of each test sequence were placed onto the consensus columns of the family CM using that sequence’s cmalign alignment. For each pair of consensus columns, these scores were then averaged over all test sequences aligned at both columns. Only nucleotides aligned to consensus columns contribute to these maps; flanking sequence and nucleotides in CM insertions are not represented. Some pairs of consensus columns had no sequence aligned at both columns, so they have no averaged score. These pairs were excluded from the ranking and are displayed as below the cut-off. Consensus columns at which no sequence was aligned were omitted from the maps. Each defined averaged score was converted to a percentile. Percentiles were computed among all defined consensus-column pairs *i*, *j* with *j i* 4, where *i* and *j* index consensus columns. The upper triangle shows pairs at or above the 95th percentile, and the lower triangle shows the consensus base pairs.

### 4.7. Calculation of token-level entropies

Token-level entropies of intergenic sequences were calculated from the logits of Evo 2 or Evo-PIN on a single forward pass of the forward and reverse strands of each padded intergenic sequence (**Fig. 1E**, **S2C**). Raw logits were converted to a categorical distribution by applying a softmax function over the vocabulary, and the Shannon entropy (*H_i_*) was calculated from the distribution as follows:

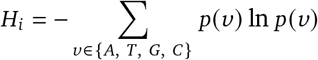

For reference, a uniform distribution over the four nucleotides (A, T, G, C) would give an entropy of ln 4 ≈ 1.39 nats. Since in autoregressive language models, the entropy calculated at position *i* is calculated for the probabilities of the next token at position *i* + 1, the entropy computed at position *i* was re-indexed to position *i* + 1, so that entry *i* of the entropy track corresponds to the base at position *i*.

To compare the token entropies of positions annotated with ncRNAs versus those that are not annotated (**Fig. 1F**), we randomly subsampled 1M intergenic loci from the 10M benchmark set. For each model (Evo 2 and Evo-PIN), the Shannon entropy was computed on both the forward and reverse-complement strand. The entropy on each position was scored as the minimum of its two strand entropies. Positions that are covered by a strict ncRNA annotation were labeled as annotated, and all other positions as unannotated. This gave 620,970 annotated positions in 4,166 loci (339 distinct ncRNA families) and 202,320,340 unannotated positions. Each annotated locus was assigned to the ncRNA family of its longest annotation, and the 995,834 loci without any annotations were randomly divided into 10,000 blocks.

For each model (Evo 2 and Evo-PIN), the change in entropy between the unannotated and annotated positions was defined as *D* = 1 *m_a_ m_u_* where *m_a_* and *m_u_* are the median entropies of the annotated and unannotated per-position entropies. We tested whether the decrease was larger for the fine-tuned model (Δ*D* = *D*_ft_ − *D*_base_) with a null hypothesis of Δ*D* = 0 with a paired permutation test. In each of 10,000 permutations, the model labels were swapped with probability ½, independently for every ncRNA family and every block, by exchanging the two models’ entropies for all positions in that unit. Δ*D* was then recomputed for each permutation. The two-sided *p*-value was calculated as (1 + number of permutations with |Δ*D*^∗^| ≥ | ΔD_observed_|)/(10,000 + 1).

For the comparison of forward and reverse strand entropies of the same position, we randomly subsampled 1M sequences from the 750M representative intergenic sequences, extracted their forward and reverse strand Evo-PIN entropies, and randomly sampled 100,000 positions from the subsampled sequences. The forward and reverse entropies were plotted for these subsampled positions (**Fig. S2D**).

To determine the entropy threshold for low-entropy tokens, we sampled the lengths of 1M padded intergenic sequences from the 750M padded intergenic sequences. For each length, we randomly generated a sequence of the same length by sampling uniformly from the four nucleotides. Evo-PIN token entropies were extracted from these random sequences using the same process described above. The forward and reverse strand entropies were concatenated, and the bottom 1% (1.3047 nats) among these entropies was selected as the low-entropy threshold (**Fig. S2E**).

### 4.8. Calculation of low-entropy pooled Evo-PIN embeddings from intergenic regions

For each padded intergenic sequence and its reverse complement, position-wise embeddings were extracted from the pre-normalization layer of every block, while token entropies were computed once from the model’s output logits as described in the previous sections (**Fig. 1G**). For each strand, we masked out the embeddings from positions in the padding sequences on either side, and positions whose entropies were equal to or above the low entropy threshold (1.3047 nats). The embeddings from the remaining positions were averaged to yield the low-entropy pooled embeddings of size D for each strand. Then, we calculated the weighted average of the two pooled embeddings from each strand, using the number of low-entropy tokens for each strand as the weights. This was done to ensure that each embedding from a low-entropy token is equally weighted in the final pooled embedding. Cluster quality scores were calculated from these low-entropy pooled embeddings extracted from each block of Evo-PIN, and compared to choose the optimal Evo-PIN block (block 25) for embedding extraction (**Fig. S2F**).

### 4.9. Clustering of pooled embeddings

Full-scale clustering was done on the low-entropy pooled Evo-PIN embeddings of the 750M intergenic sequences while excluding three categories of sequences. First, we removed 128M intergenic sequences derived from the edges of metagenomic contigs that are bounded by a contig boundary, as the difficulty of gene calling near contig ends meant these regions may contain unannotated protein-coding sequences. Second, we removed 1.5M sequences containing ambiguous (non-A/T/G/C) bases. Third, we removed 223M sequences with fewer than 10 low-entropy tokens on each of the forward and reverse strands, as these pooled embeddings were likely to be more noisy from a smaller sample size and these sequences were unlikely to contain ncRNAs, as 99.8% of sequences with known ncRNA annotations contained 10 or more low-entropy tokens (**Fig. S3F**).

To cluster the remaining 399M low-entropy pooled embeddings, we first constructed an approximate k-nearest-neighbor graph (*k* = 16) using algorithms from the Faiss library (61). For a more efficient neighbor search, we used an inverted file index (62) with product quantization (63) (IVFPQ), which partitioned the embedding space into Voronoi cells and compressed the vectors within e_√_a<u>ch</u> cell. The index was trained on 4M randomly sampled embeddings with 79,859 centroids (chosen to be 4 *N*, with N = 399M) and 512 8-bit sub-quantizers with an optimized product quantization (OPQ) rotation (64), and was used to compress each embedding from 16 kilobytes to 512 bytes. Centroid lookup was accelerated using the Hierarchical Navigable Small World (HNSW) graph (65) (m=32, efConstruction=40). At query time, 128 cells were probed per query (HNSW efSearch=64). The resulting k-nearest-neighbor graph was partitioned using the Leiden algorithm (66) (CPM/modularity) implemented by NetworKit (67) v11.2.1 with a range of resolution parameters.

### 4.10. Ablation experiments for clustering

We compared the end-to-end query throughput for an inverted file index with product quantization (IVFPQ) and a Hierarchical navigable small world (HNSW) coarse quantizer with a flat (brute-force)_√_i<u>n</u>dex at database sizes of 500K, 1M, 2M, 5M, and 10M embeddings. The IVFPQ index used PQ512 with 4 *N* centroids, and nprobe was set to 0.2% of nlist. Throughput was measured in queries per second (**Fig. S3A**).

To evaluate the effect of vector compression on the cluster quality scores, we sampled 1M sequences as a training set to construct the indices with various combinations of quantizer configurations – namely three sub-quantizer settings (PQ128, PQ256, PQ5_√_1<u>2</u>), eac_√_h<u>w</u>ith and without OPQ rotation. The number of inverted file index (IVF) centroids was set to either 4 *N* or 8 *N*, where N was the number of sequences annotated with ncRNAs. All queries were searched with nprobe set to be equal to nlist (i.e., exhaustive cell search). Cluster quality scores were evaluated on the same subset of 10M intergenic sequences that was used to benchmark the Evo 2 fine-tuning and pooling functions (**Fig. S3B**).

We characterized the tradeoff between search accuracy and the number of cells searched at query time, by training a PQ512 index on 1M embeddings a_√_nd building indices at varying database sizes (100k, 500k, 1M, 2M, 5M and 10M), while scaling nlist at 4 *N*. For each configuration, nprobe was swept as a fraction of nlist (0.1%, 0.2%, 0.5%, 1%, 2%, 5% and 10%) to ensure comparable coverage across database sizes. A fixed set of 10k query embeddings was used across all conditions and recall was measured against a flat (exact) index (**Fig. S3C**).

We also benchmarked the end-to-end query throughput for the IVFPQ index with and without a HNSW coarse quantizer at database sizes of 500K, 1M, 2M, 5M, 10M, 20M and 50M embeddings for all indices (**Fig. S3D**). To investigate the speed-quality tradeoff of a coarse quantizer, we compared the recall of both indices against a brute force search when searching an index of 10M embeddings (**Fig. S3E**).

### 4.11. UMAP visualization of pooled embeddings

We visualized 32,000,540 sequences from 255,213 clusters that were a union of these four groups of clusters: (i) 85,707 clusters whose CMfinder motifs had strong covariation support and one or more significantly colocalized genes (see **Structural classification of representative CMfinder motifs** and **Identifying colocalized protein annotations**), (ii) 122,127 clusters whose CMfinder motifs had strong covariation support with at least 2 separate hairpin structures each with at least 2 significantly covarying base pairs, and had no significantly colocalized genes, (iii) 46,148 clusters whose CMfinder motifs had weak covariation support, but had strong colocalization signals ( *f_c_* > 0.5), and (iv) 1,672 clusters with periodic token entropy patterns (see **Searching for clusters with periodic entropy patterns**). This same subset of clusters was also selected for display through the web app. The two-dimensional UMAP embedding was used to map each sequence to a 2D coordinate (umap-learn v0.5.11) (68). To avoid recomputing nearest neighbors, we restricted the previously computed nearest-neighbor graph to this subset of sequences, initialized edges as the cosine distances, and replaced the edges to sequences outside of the subset with edges to itself with distance 0. We then used the umap-learn functions fuzzy_simplicial_set to construct the fuzzy simplicial set and simplicial_set_embedding to optimize its two-dimensional layout for 400 epochs with min_dist = 0.1.

### 4.12. Selection of the Leiden resolution parameter

The Leiden resolution was chosen by optimizing for the number of known ncRNA families that are grouped into high-purity clusters of size 50 or larger. We experimented with 12 Leiden resolution values (10^2^, 2 10^2^, 5 10^2^, 10^3^, 2 10^3^, 5 10^3^, 10^4^, 2 10^4^, 5 10^4^, 10^5^, 2 10^5^, 5 10^5^) for the clustering of the pooled embeddings. After clustering the pooled embeddings at each resolution, the sequences within each cluster were deduplicated again at 70% sequence identity using MMseqs2 (56) to remove duplicate sequences that escaped detection at the full set of 750M sequences, which lowered the total sequence counts across all clusters from 399M to 376M. We observed that as the resolution increased, the number of clusters increased, while the median cluster size (the number of deduplicated sequences in each cluster) excluding singletons increased until resolution 103, and then started decreasing monotonically (**Fig. S3G**).

The objective of the clustering was to cleanly split intergenic sequences with distinct noncoding elements while keeping the cluster sizes large enough such that the downstream analyses have enough statistical power. Thus, for each resolution, we selected for clusters that (i) had at least 50 deduplicated sequences in total, (ii) had at least 10 sequences uniquely annotated with a known ncRNA according to the strict annotations, and (iii) had an annotation consistency (the fraction of annotated sequences that were annotated with the most common ncRNA in the cluster) of at least 0.9. Within these clusters, we counted the total number of known ncRNAs that are represented as the most common ncRNA. Choosing the Leiden resolution of 10^5^ maximized this count at 676 families. Increasing the resolution further increased the number of unique ncRNA families being split into high-consistency clusters, but these clusters were often split into clusters smaller than size 50.

### 4.13. Calculation of median pairwise sequence identities of sequences within each cluster

To calculate the pairwise sequence identities of the sequences within each cluster, we first took a random subsample of up to 50 deduplicated sequences from each cluster. The subsampled sequences of each cluster were aligned with MAFFT v7.490 (69) using the FFT-NS-2 progressive algorithm (--retree 2), with the --adjustdirection option enabled such that each sequence was reverse complemented when it improved the alignment. For every pair of sequences in the alignment, Easel esl-alipid (v0.49, DNA mode; https://github.com/EddyRivasLab/easel) distributed with Infernal v1.1.5 (58) was used to calculate the percent identity as the number of identical positions divided by the ungapped length of the shorter sequence. The median of these pairwise identity values was reported for each cluster (**Fig. S4B**). 759 singleton clusters were skipped as no pair could be formed.

### 4.14. Embedding-based reorientation of sequences within each cluster

For each sequence *i* within a cluster, we started from their low-entropy pooled Evo-PIN embeddings from their forward strand ( *f_i_*), and their reverse strand (*r_i_*). Both pooled embeddings were L2-normalized. The goal of the reorientation algorithm was to look for a binary flip vector *s* 0, 1 *^n^* for each cluster, where *s_i_* = 1 means that sequence *i* was reverse complemented from its original orientation and *n* is the number of sequences in the cluster. All *s_i_* were set to 0 initially.

For each iteration of the algorithm, the following steps were performed:

i. Compute the cluster means *μ _f_* and *μ_r_* for the forward and reverse pooled embeddings
ii. Determine whether each sequence’s forward and reverse pooled embeddings are closer to the cluster’s respective means compared to the other mean (i.e. compare dot product *f_i_* · *μ _f_* + *r_i_* · *μ_r_* against *r_i_* · *μ _f_* + *f_i_* · *μ_r_*)
iii. Toggle *s_i_* for every sequence where flipping brings the *f_i_* and *r_i_* closer to their cluster means, updating all sequences at once

The loop terminated when no *s_i_* within a cluster changed in an iteration. Once *s* had converged, we flipped the values of all *s_i_* in a cluster if more genes that are adjacent to the intergenic sequences of the cluster were on the reverse strand, such that more adjacent genes would always be placed on the forward strand across the cluster. This was done to maximize the probability that the shared noncoding element is placed on the forward strand of each intergenic sequence, under the assumption that these noncoding elements would more likely be on the same strand with their neighboring genes.

To validate the results of this algorithm, we used the clusters with at least 3 sequences that share the annotation for the same ncRNA using the strict annotations (n = 28,223), and checked the strandedness of the known ncRNA calls before and after the reorientation. We confirmed that for these clusters with known ncRNAs, this algorithm achieves perfect strand consistency for 80% of the clusters, and >90% consistency for 91.4% of the clusters (**Fig. S6**).

### 4.15. Identification of shared noncoding motifs with CMfinder

From each cluster with at least 3 sequences, we randomly subsampled up to 20 sequences and used CMfinder\ 0.4.1.9 (cmfinder04.pl, with Infernal 1.1.5) (25) to find conserved structured motifs. We used the options -combine -minCandScoreInFinal 10 -cpu 1, plus the expectation-maximization options --filter-non-frag --fragmentary --max-degen-per-hit 2 --max-degen-flanking-nucs 7 --degen-keep --amaa. ClustalW-based merging of motifs across gaps was turned on such that CMfinder could join single-hairpin and double-hairpin motifs into larger combined motifs. The initial CMfinder run on all clusters returned 11,531,700 motifs in 2,675,408 clusters.

We first discarded motifs that were built from seed sequences that covered less than 75% of the input sequences to CMfinder. Because CMfinder emits many merge-depth variants of the same structure, we reduced each cluster’s motifs to a non-redundant set using the following filters. First, a motif was removed if its components were a strict subset of another surviving motif’s components (for example, h1_3 when h1_3.h2_5 also survived). Second, a motif was removed if at least 90% of the positions that the motif covers are within the union of the coverage of the other motifs still remaining, while processing the motifs from smallest to largest in number of positions covered throughout the cluster. For clusters that did not have any motifs that survived these filters, we reran the same filters with the initial seed coverage threshold decreased to 50%.

Each remaining motif alignment at this stage was converted to a covariance model with cmbuild. Each model was searched against all deduplicated, reoriented sequences in its cluster with cmsearch –max --toponly -T 15. The best forward-strand hit per sequence was aligned to the model with cmalign. If the sequences used as input to CMfinder already included every deduplicated member, the CMfinder alignment was used directly. Each alignment at this stage that was extended to the full cluster was scored with R-scape 2.0.4 (30) using CMfinder’s predicted structure (--gapthresh 0.5). Motifs with no base pairs left after removing alignment columns that were more than 50% gaps were discarded. After this step, 3,317,848 motifs remained in 1,952,307 clusters.

We then chose a single motif per cluster to be the cluster’s representative motif by three rules, applied in order: (i) most base pairs in the predicted structure; (ii) if tied, the longest consensus structure (total SS_cons columns in the extended alignment); (iii) if still tied, the first motif file name in alphabetical order. Clusters were kept for downstream motif classification if their representative motif, after the extension to the full cluster, was found in at least 50% of the cluster’s deduplicated sequences. This process resulted in 1,516,886 clusters with a single representative motif each.

### 4.16. Structural classification of representative CMfinder motifs

Each of the representative CMfinder motifs of the 1,516,886 Leiden clusters was classified into the four structural categories using the following steps. First, we used TerminatorNet (29) to annotate the locations of all rho-independent intrinsic terminators on every member sequence of the 1,516,886 clusters. Terminator annotations with score 0.9 were kept. A cluster’s representative motif was labeled a likely terminator if all of the following conditions were true: (i) its consensus secondary structure was a single hairpin with no pseudoknot; (ii) a TerminatorNet terminator annotation overlapped the motif in ≥10% of member sequences, and in at least 2 sequences; and (iii) in ≥80% of the cluster’s member sequences, the flanking protein-coding gene upstream of the terminator annotation was on the same strand as the terminator. These conditions ensured that motifs were labeled likely terminators only if the structural and contextual evidence both agree for the given motif.

All remaining motifs were classified by the covariation statistics of the RNA structure predicted by CaCoFold (31). CaCoFold-predicted structures and covariation statistics were calculated by passing the motif’s alignment through R-scape with the CaCoFold option, while keeping alignment columns with less than 50% gaps (R-scape --cacofold --gapthresh 0.5 --covmin 4 --minhloop 3). From these structures, we parsed the total number of base pairs in the structure, the number of significantly covarying pairs among those base pairs (e-value < 0.05), and the expected number of significantly covarying base pairs i.e. the sum of R-scape’s per-pair statistical power. This was the number of covarying pairs that are expected to be detected given how much variation the sequences in the alignment contain.

A motif was classified as having no covariation support if the number of significantly covarying base pairs was zero, or the expected number of significantly covarying base pairs divided by the total number of base pairs (i.e. the power of the alignment) was less than 0.10, indicating not enough variation existed in the aligned sequences to test covariation (70). A motif was classified as having strong covariation support if the motif’s predicted structure had at least 5 significantly covarying base pairs, and the ratio of the observed number of significantly covarying pairs to the expected number was at least 0.50. All remaining motifs were classified as having weak covariance support.

### 4.17. Identifying colocalized protein annotations

For each intergenic region, we recorded the amino acid sequences of every protein-coding gene called on the same contig within 5 kb of the locus. A gene counted as neighboring if its end was no more than 5,000 bp upstream of the locus start and its start was no more than 5,000 bp downstream of the locus end. This resulted in 1,471,319,620 protein sequences in total, which were clustered with MMseqs2 linclust (v18-8cc5c) (56) at 90% sequence identity and 80% alignment coverage to 413,734,648 cluster representatives.

We then annotated the representative amino acid sequences with InterProScan 5.75-106.0 (71), using the Pfam member database (-appl Pfam, Pfam 37.4) (72). The full InterProScan run produced 168.2M Pfam domain hits across 123,809,676 representative proteins. For each representative, we sorted its Pfam hits by their start positions on the protein. We joined the accession IDs of the Pfam hits in that order into an N-to-C-terminal domain architecture (e.g. PF02838|PF00728). Every member of an MMseqs2 cluster then inherited its representative’s architecture. Proteins whose representative had no hits were given an empty domain architecture. After this step, 922,183,617 of the 1,471,319,620 proteins (62.7%) had at least one Pfam domain annotation: 78.4% of proteins in GTDB genomes and 56.0% of those from metagenomes.

To identify protein domain architectures that colocalize significantly with the intergenic sequences of each cluster, we counted the total number of times a domain architecture is found adjacent to or nearby (within 5 kb) each deduplicated locus within a cluster. For each (cluster, architecture) pair, we then ran a one-sided Fisher’s exact test (hypergeometric upper tail) (73) using SciPy 1.16.1 (74) against a background of every flanking gene across all clusters (N = 752,010,400) and every nearby gene across all clusters (N = 2,868,784,646). A separate enrichment *p*-value was calculated for flanking and nearby architectures, and each was Bonferroni-corrected for every pair tested (355,388,431 pairs for adjacent genes, 1,296,530,951 pairs for nearby genes). Protein architectures were marked as significantly colocalized in the flanks of a cluster if the corrected *p*-value ≤ 10^−30^ and at least 20% of the sequences in the cluster flanked the protein architecture. Likewise, protein architectures were marked as significantly colocalized near a cluster if the respective corrected *p*-value ≤ 10^−30^ and at least 20% of the sequences in the cluster were found near the protein architecture.

### 4.18. Identifying colocalized protein folds

All amino acid sequences of the 413,734,648 90% identity cluster representatives of proteins within 5 kb of the intergenic loci were translated into the Foldseek 3Di-tokens using the ProstT5 model (encoder-only 3Di prediction) (75). We then used Foldseek to cluster the collected proteins for each cluster individually, with coverage set to 0.6, an e-value cutoff of 1e-8, and the minimum sequence identity set to 0 (foldseek cluster --min-seq-id 0 -c 0.6 -e 1e-8) (5). We left the alignment type at the default setting requiring both query and target to pass the coverage threshold and searched only the 3Di and amino acid sequences since the ProstT5 predictions did not return any coordinates. The representative sequence of each cluster that contains more than 1 protein was then searched against the PDB100 (snapshot 250101) (76) and Swiss-Prot (v6, March 2025) (77) databases clustered at coverage 0.9 with an e-value cutoff of 1e-10, which reduced the database size from 324,204 to 97,463 members for PDB and 590,183 to 94,744 members for Swiss-Prot.

### 4.19. Searching for clusters with periodic entropy patterns

For each intergenic sequence, we took the position-wise minimum between the forward strand token entropies and reverse strand entropies of Evo-PIN, and mean-centered the entropies to build a single token entropy track across the sequence. Sequences shorter than 30 nt, and sequences whose entropies had no variation (s.d. < 10^−6^) were excluded. The autocorrelation of each entropy track was computed with a zero-padded Fast Fourier Transform available in NumPy 2.3.2 (78) via its pocketfft backend for lag *p* values ranging from 0 to 120 nt, and normalized to 1 at *p* = 0. The autocorrelation values of all usable sequences in a cluster were averaged for each lag value. Clusters with fewer than 20 usable sequences were not scored.

For each cluster, we looked for local maxima of the mean autocorrelation between lags *p* = 2 and *p* = 120 using SciPy’s find_peaks function (v1.16.1) (74), and computed each peak’s topographic prominence. The periodicity score was set as the prominence of the most prominent peak, and the dominant period *p* was that peak’s lag. Since protein-coding sequences that are missed by Prodigal (53) would produce a strong 3-nt entropy period from every codon’s third-position (wobble) uncertainty, we removed any clusters whose dominant period was 3 nt, or clusters whose dominant period was a multiple of 3 and whose mean autocorrelation also had a peak at lag 3. Of the 5,138,521 clusters, 4,951,832 clusters had at least 20 usable sequences and were scored. We called a cluster as having periodic entropy patterns if its periodicity score was ≥ 0.7, giving 1,672 periodic clusters (**Fig. 3A**).

### 4.20. Structural comparison of context-enriched proteins to known array-associated proteins

For each Leiden cluster with periodic entropy patterns, we identified the largest Foldseek cluster (5) among its neighboring proteins, and folded the representative sequence from that cluster using ESMFold (4). For the sequences of known array-associated proteins, we used the PDB ID: 8J12 (retrieved June 2026) (76) for Cas12f, UniProt ID: X1U2U4 for the Vipr protein reported in its original discovery (16) and the protein sequence reported in the TIGR-Tas discovery manuscript (18) for FpTasA. We used ESMFold to predict the structures of these reference proteins. We aligned the structures of the target and reference proteins using Foldseek’s easy-search with alignment-mode=1 (TM-align). Given the aligned C*α* coordinates, we then superimposed the two predicted structures using the Superimposer class implemented in the biopython PDB module (version 1.82) (79). The superimposed structures were visualized using ChimeraX (version 1.12-rc2026.05.11) (80).

### 4.21. Analysis and typing of CRISPR-like arrays

We selected 1,481 clusters (317,021 deduplicated sequences in total) with periodic entropy patterns whose dominant period *p* lay within the 60-79 nt band. For each cluster, we used its deduplicated, reoriented member sequences as an input to MinCED v0.4.2 (34), which implements the CRISPR Recognition Tool (CRT) algorithm (81). MinCED was run with a 900s timeout and the options -minNR 2 -minRL 18 -maxRL 55 -minSL 18 -maxSL 60 -spacers, from which 292,368 (92.2%) of the sequences returned an output. From MinCED’s output, we parsed the array coordinates, number of repeat copies, and the mean repeat and spacer lengths. When a locus carried more than one array, its dominant array was chosen as the one with the most repeat copies. This array’s mean repeat and spacer lengths were set as the locus’s repeat and spacer lengths (**Fig. S9A**). The sequences of the repeats were parsed from the dominant array, keeping only the repeats with the most common repeat length for that array. The most common base in each position of these repeats was used for the consensus repeat sequence for each locus.

To measure the consistencies of the repeat/spacer lengths and repeat sequences within each cluster and between clusters, we randomly selected 100 loci per cluster, with replacement, so that every cluster contributed equally regardless of size. Each selected locus was paired with (i) a different locus from the same cluster (within-cluster pair) and (ii) a locus from any other cluster, drawn from all loci (between-cluster pair). The two sequences in a pair were required to originate from different sources (different genomes for GTDB-sourced sequences, different samples for metagenomic sequences). This was done to prevent several arrays from one genome or metagenomic sample inflating the within-cluster similarity. A pairing that failed this rule was redrawn up to 20 times, after which the locus was dropped. In total, we compared the spacer/repeat lengths for 147,790 within-cluster and 148,100 between-cluster pairs, and the repeat sequences for 147,802 within-cluster and 148,100 between-cluster pairs. For the repeat/spacer length difference, we used the absolute difference between the two loci’s repeat/spacer lengths. For the direct repeat’s sequence identity, we measured the best-matching gap-free identity between the two repeat sequences by sliding one repeat across the other and dividing the number of matching bases by the length of the longer repeat (**Fig. S9B**).

Each sequence within the 1,481 clusters was also independently typed with the CRISPRCasTyper classification tool (cctyper v1.8.0) (35). For each locus, we extracted the amino acid sequences of all proteins within ±20 kb of the loci, and searched these sequences against the 705 Cas profile HMMs distributed with cctyper v1.8.0 using HMMER v3.4 (hmmsearch --domtblout --noali -E 1e-3). For each protein, the highest-scoring domain hit was retained then filtered such that profiles belonging to a family with a class-2-specific cutoff in cutoffs.tab were required to meet that family’s e-value, sequence coverage and HMM coverage thresholds, and all remaining profiles were required to satisfy the default thresholds (*E* < 0.01, sequence coverage 0.3, HMM coverage ≥ 0.3). Retained hits were grouped into operons using cctyper’s adjacency rule, with cas genes separated by at most 3 unannotated genes assigned to the same operon. Each operon was then typed by using cctyper’s type_operon() routine (cctyper/castyping.py, v1.8.0). If a locus contained more than one typed operon, the highest-scoring operon was taken as the call for the locus. 96,083 (30.3%) loci across 1,352 clusters had a type call. The type/subtype consistency was plotted only for the clusters with at least 5 typed loci (**Fig. S9C**).

### 4.22. Identifying repeat arrays in VIPR array-like clusters

From the 99 clusters with dominant periods of 5, we first selected 76 clusters whose sequences colocalized with a gene encoding for a protein similar to the reported Vipr protein (16). For each cluster, the structures of the representative proteins of its five largest Foldseek clusters were predicted with ESMFold (4). Each prediction was aligned to the AlphaFold DB model of X1U2U4 with Foldseek (alignment-mode=1, TM-align). We kept clusters whose best representative protein had a mean pLDDT of at least 0.6 and was aligned to at least 120 residues of X1U2U4.

For the sequences in these 76 clusters, repeat units were modeled as 5 nt patterns of the form XYZNN: three constrained positions followed by two unconstrained ones. Each of X, Y and Z was set to be any of the 15 IUPAC nucleotide codes, giving 3,375 candidate cores. Each core was searched across the deduplicated, reoriented sequences of every cluster. A 5-mer matched the core only if every base was allowed at each position with no mismatches. Tandem arrays were called independently in each of the five frames as runs of ≥3 adjacent matching units that are exactly 5 nt apart. Arrays lying entirely inside a longer array were removed. For each cluster-core pair, we recorded the total number of repeat units in arrays, and each core was scored by its excess repeat units over a shuffled null, weighted by information content. The null sequences came from five mononucleotide-shuffled copies per every sequence in the cluster. The information content (IC) of each 5-mer was computed using the cluster’s own base composition as follows:

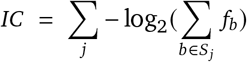

Here, *S_j_* is the set of bases allowed at position *j* and *f_b_* is the frequency of base *b* among the sequences in the cluster. By design, a position allowing all four bases contributed nothing to the information content. The score for each core sequence was calculated as follows:

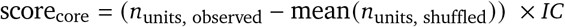

For each cluster, the core was initially set to the highest-scoring of the 64 concrete trinucleotides without degenerate bases. It was then relaxed greedily – the score of every pattern obtained by adding one degenerate base to the allowed set at a single position (concrete to two-fold code, two-fold to three-fold, three-fold to N) was compared to the current score. The best of these new relaxed patterns was accepted only if it improved the score by ≥50% relative to the current score, and the process repeated until no single step met this threshold. The final relaxed core was what was used as the cluster’s fixed 3-mer (**Fig. S10A**).

We used each cluster’s core 3-mer to find every exact match within the sequences of the cluster to the core. A core match with one mismatch was also accepted, but only if it lay exactly 5 nt upstream or downstream of an exact match. Found units were kept only when they formed runs of at least three, spaced exactly 5 nt apart. Where runs in different frames overlapped, units from the longest run were kept first, and any unit starting within 5 nt of an already kept unit was discarded. Each cluster’s sequence logo of the 5-mer was built from all repeat units found across the sequences of the cluster using the rules above. Units extending past the end of a sequence or containing an ambiguous base were excluded. The tandem motifs from the 10 longest loci from each cluster with noncanonical 3-mers were plotted (**Fig. S10B**).

### 4.23. Clustering of intergenic sequences adjacent to IS110 recombinases

We selected 2,332 clusters where the Pfam domain architecture of the most common significantly colocalized flanking protein was identical to the domain architecture of the IS110 recombinase – the DEDD_Tnp_IS110 (PF01548) domain followed by a Transposase_20 (PF02371). From the 405,556 deduplicated sequences of these clusters, we removed any sequences that are 50 nt or shorter, or those without the recombinase as their flanking protein. The remaining 181,180 sequences were clustered following the same approach taken for the global clustering, with the exception of an exact nearest neighbor calculation replacing the approximate index used for the full set. The Leiden resolution was set to 10.

### 4.24. Construction of cluster-specific covariance models of bridge RNAs

For each cluster with at least 3 intergenic loci, we randomly selected 100 loci, or the total number of loci per cluster, whichever was smaller. The loci were then aligned using MAFFT Q-INSI (MAFFT 7.525) (69). Every column in which ≥50% of sequences have a gap was dropped and sequences that were empty after this gap filter were removed. An initial consensus structure was predicted from this alignment using R-scape CaCoFold (R-scape --cacofold --gapthresh 0.5, R-scape 2.0.4a.) (30, 31).

Next, we trimmed the unaligned columns back to three columns beyond the first and last unaligned columns. We then built a covariance model (CM) from the seed alignment (Infernal v1.1.5), calibrated the model, and searched each CM against the loci in the cluster it was created from. For refinement, we took the best forward strand hit of every locus at *E* ≤ 10^−3^ (top 2,000 by bitscore) and aligned those hit sequences to the model using cmalign. We rebuilt the model from that alignment, keeping the seed’s consensus structure (cmbuild --hand). Models with fewer than five hits in the search round were discarded.

The refined models were then again trimmed by removing the tails without base pairing on both sides. Then, working inwards from the 5^′^ and 3^′^ ends, we trimmed all hairpins smaller than 12 base pairs that did not have at least 4 base pairs with one of them showing significant covariation (R-scape e-value ≤ 0.05), stopping at the first domain exceeding those criteria. 3 nt of padding were retained on the start and end of the first and last retained domain respectively. If no domain with at least one covarying base pair could be found, only the unpaired edges were trimmed. The trimmed models were then refined once more by searching for hits in the cluster as described in the previous section. Only models that hit more than 50% of loci in their own clusters with an e-value of ≤ 10^−3^ were retained.

All retained CMs were then fully calibrated and searched against the full database (cmsearch -Z 131.802782) of IS110-adjacent intergenic sequences. All locus-CM pair hits were retained if their e-value was ≤ 10^−3^ and their bitscore was over 50.

### 4.25. Reconstructing target and donor sites of candidate bridge RNAs

Boundaries of IS110 elements were derived as previously described (14). In brief, we extended the IS110 recombinase CDS by 1,000 nt on each side and searched 100 nt fragments on each side against the metagenome database as well as an extended version of the GTDB r226 database where we randomly selected up to 10 non-representative genome assemblies per representative genome. If the two fragments formed a consecutive stretch in a contig we retained it as a candidate empty site. Boundaries were then refined using a modified version of the MGEfinder “wholegenome” command (82) published previously (14). We then expanded the initial set of elements by iteratively searching the now verified boundaries of elements against all padded recombinase loci using blastn. We accepted the predicted boundaries of a new element, if the 5^′^ and 3^′^ ends of another element with verified boundaries were matching, the length of that new hypothetical element was within ±5% of the original, and the boundaries matched the predicted core for the reference element, in case MGEfinder reported a core. These new hits were then added to the query set. This search process was repeated a total of 16 times.

After these iterations, a final pass searched all records without assigned boundaries using up to 50 elements per cluster as queries. A second set of elements were kept if they were within ±10% of the cluster mode of element length and the terminal 10 nt aligned within 15 nt of the end of the blast hit, with at most 2 mismatches.

From each of those boundary calls, we reconstructed 50 nt long target and donor sites for each element. Target sites were constructed as the concatenation of the 25 nt sequence 5^′^ of the core, the core of length *c*, and 25 *c* nt sequence after the core in the pre-insertion site. For the donor, we concatenated the last 25 nt of the element with the core and the first 25 *c* nt of the element, representing the sequence of the excised element around the shared core.

For elements without a predicted core, we retrieved the 4 most common predicted cores in other elements hit by the same CM, then searched the region flanking the preliminary element boundaries. If any of the cores flanked both ends of the elements at the same shift of at most ±1 nt, we used that core and the new element boundaries. All elements where no precise boundaries could be inferred were discarded from the following analyses.

### 4.26. Covariation analysis between bridge RNAs and donor/target sites

We first searched all IS110 elements with predicted boundaries with the CMs corresponding to their intergenic loci in the previous search steps. All hits covering at least 80% of the model and overlapping with the predicted element were retained. We then aligned the hits using cmalign (--nonbanded, --notrunc) and removed all residues falling outside of each element’s predicted boundaries. All residues overlapping with the 25 nt windows used to reconstruct the donor site were also removed.

For each CM, we then retrieved the reconstructed target and donor sites, and used MMseqs2 (mmseqs easy-linclust, 95% identity, 95% coverage) (56) to cluster the rows (without the gaps) of the aligned CM hits, targets, and donors, respectively. One row was kept for each CM cluster x target/donor cluster pair, preferring rows with cores that were called directly over rescued core assignments. If an alignment had fewer than 20 such rows it was not analyzed further.

The following covariation analysis follows the previously described method to discover covarying bases between bridge RNAs and target or donor sites (14): we concatenated the CM alignment with the target and donor sequences separately, which were aligned by centering on the first core position. Then, we analyzed the alignments for covariation using CCMpred (100 iterations) (83). In addition, we calculated base pairing concordance scores as previously described, which give positive values to covarying nucleotides that form Watson–Crick base pairs on the forward strand while assigning negative values to positions that are more likely to pair with the reverse strand. We then min-max scaled the covariation matrix and negated any value with a negative concordance score.

### 4.27. Prediction of potential guide lengths

For all CMs, we manually inspected the covariance between the RNA and the target and donor sequences along with the consensus structure, and manually selected 86 models with clear covariation on 4 non-overlapping sites that lined up with plausible positions in the predicted secondary structure. Within this curated set, we then assigned covarying regions by selecting cells from the covariation matrix whose absolute value post normalization is in the top 1% for its covariance matrix. Any potential runs of two or more such cells with the highest cell required to have an absolute value >= 0.4 lined up along a diagonal or antidiagonal streak, with gaps of up to 2 cells, and all cells having the same sign in the covariance matrix, were marked. Potential runs were then filtered to only those overlapping with the ±10 nt region around the predicted core.

We then collected all valid combinations of runs that satisfy the following criteria: (i) There should be two called runs in the covariation map between the RNA and the target sequence as well as two runs of covariation detected in the RNA to donor covariation map. (ii) Of the two covariation runs assigned to the target or donor region one has to have a negative concordance score, while the other has to have a positive score. (iii) No called site should overlap with any other site in the combination.

We then iterated over all combinations and searched them for potential base pairing between RNA and DNA using pairwise alignments. We extracted the DNA with a padding of 5 nts on each side around the covariation run and RNA sequence with a padding of 7 nts. If the run was antidiagonal, we reversed the RNA sequence, and for runs with a negative coupling value we reverse-complemented the DNA sequence. We then ran an ungapped, banded local alignment on those residues scoring +1 for each matching pair and −2.5 per mismatch within a band of 2 around the identified coupling run. We retained the highest scoring alignment overlapping with the covariation run on the DNA side. We retained the combination that has the highest total number of potential pairs summed over all 4 separate sides and at least 3 nt matching per site. We then generate a null value for the used combination of sites by calculating the predicted base pairing to a randomly selected RNA that is assigned to the same CM.

### 4.28. Visualization of bridge RNA Leiden clusters

We computed a centroid for every Leiden cluster that contains at least one locus with a TPN estimate (220 clusters). The centroid is the mean of the L2-normalized embeddings of all intact members of the cluster (102,948 loci). Centroids were joined by average linkage (UPGMA) on Euclidean distances, using SciPy (v1.18.1) with optimal leaf ordering. Each TPN-scored locus (n = 32,190) was attached as a leaf under its own Leiden cluster. Leaves hang from the cluster node on a fixed branch of 0.15 × the root height, in a seeded random order. The tree was drawn as a circular fan (ggtree, v3.10.0) with branches colored by Leiden cluster. An outer ring of radial bars shows each locus’s TPN on a fixed axis of 20-40 nt: loci below 20 sit at the baseline (2,709 loci) and none exceed 40.

### 4.29. Visualization of consensus bridge RNA structures

For each bridge RNA CM, we aligned all strict hits with a bitscore 50 using cmalign (Infernal 1.1.5); only the CM’s match columns were kept. We predicted a consensus secondary structure on this alignment with ConsAlifold (gamma = 2) (84). ConsAlifold was run on a random sample of 150 sequences. Covariation was then tested on the full alignment, not the sample using R-scape 2.0.4.a in two-set mode (-s), which tests the predicted base pairs against all other column pairs, with no CaCoFold refinement. Pairs with *e* ≤ 0.05 were called significantly covarying. Structures were visualized with R2R as bundled with R-scape.

### 4.30. Structure prediction of the bridge recombinase synaptic complex

The synaptic complex was modeled with Protenix (39), an open reimplementation of the AlphaFold3 architecture (85), using the protenix_base_20250630_v1.0.0 checkpoint. Each construct was specified as a single multi-chain assembly of four recombinase protomers (395 aa each), one bridge RNA (bRNA; 166 nt) and eight DNA oligonucleotides (173 nt in total). Each half-site duplex was supplied as four separate chains rather than two – the 5^′^ and 3^′^ segments of both strands (30/14/17/27 nt for the target site and 28/15/15/27 nt for the donor). This split was necessary because co-folding methods of this class model base-paired duplex geometry and do not perform strand separation: the unwinding and melting of the target and donor duplexes that bridge recombination requires cannot emerge from the prediction itself, so the open state has to be encoded in the input rather than generated. Protein and RNA multiple sequence alignments were precomputed and supplied directly (2,322 and 410 sequences, respectively), giving an effective alignment depth of *N_msa_* = 2,321 over 1,919 tokens and 19,313 atoms. No structural templates and no atom-contact, contact or pocket constraints were used, so that the predicted bRNA–DNA geometry was independent of the designed base pairing against which it was subsequently scored. Inference used 10 Pairformer recycles, 200 diffusion steps and bf16 precision, generating five samples per random seed on a single NVIDIA H100 80 GB GPU. The model selected (seed 2065, sample 0) formed 35 called guide pairs at a median heavy-atom distance of 2.75 Å, retained an intact bRNA backbone (largest C1^′^–C1^′^ step 9.88 Å), and placed both half-site dimers at 29.4 Å center-to-center with 990 and 1,002 atom-pair contacts, compared with 31.0 Å and 1,284 contacts in the reference IS110 synaptic complex model. Its confidence metrics were ipTM 0.859, pTM 0.879 and mean pLDDT 82.3, with no steric clashes detected.

### 4.31. Harness adaptation for agentic mining

We investigated the noncoding element clusters using compass-agent, a harness built on OpenAI Codex CLI. It configures a lead agent (gpt-6-astra, high reasoning effort) through Codex’s project-level configuration. A system level prompt defined the lead orchestrator role. These system level prompts ensured that every reported number must come from a query run in that session, with its denominator and starting cluster set stated. Furthermore, the agent was instructed that no cluster of noncoding elements may be called novel until it has been checked against the known ncRNA annotations in the cluster database based on the permissive Rfam hits, and terminator and periodic entropy pattern classifications have been ruled out.

The researcher was able to directly address the lead agent through the Codex CLI tool. The lead could split independent tasks across up to six parallel miner sub-agents (gpt-6.1-sol). Each miner worked in its own subfolder. Miners messaged siblings about findings across tasks and returned a tabular candidate list. The lead treated these lists as claims and spot-checked them before reporting. All communication between lead agent, miner sub agents and researcher was recorded.

### 4.32. Execution of unattended agent campaigns

For unattended campaigns, a supervisor agent ran as a lead orchestrator and was supplied a user-specified rubric. Each researcher-created criterion was to be marked by agents between 0 - 4. Miners were required to keep a record for every considered candidate, including rejected and known comparisons. After discovery, a separate grader agent, without the discovery conversation in context, re-ran the decisive checks and filled in the rubric evidence. These checks covered expanded alignments and covariation power, flanking-protein identities, Rfam overlap, RNA placement and primary-literature passages. Each candidate then received a final weighted score across all criteria in which missing criteria counted as zero. Retention required passing certain strict checks e.g. novelty. Independent replicate rollouts were run as SLURM job arrays, and recurrence across rollouts was reported.

### 4.33. Agent access to the dataset and analysis tools

Agents queried the dataset through a read-only Model Context Protocol (MCP) server exposing twelve typed tools for each cluster, including the cluster’s basic properties, cluster membership, sequence files, CMfinder motifs, and annotations of proteins or shared folds in the genomic contexts. The same server exposed a restricted SQL interface that accepts only read statements, rejects any statement containing data-definition or data-manipulation verbs, and caps query memory at 6 GB. A second MCP server provided literature search against Europe PMC and PubMed. Agents additionally had shell access to the analysis binaries, including those used elsewhere in this work (Infernal 1.1.5, CMfinder 0.4.1.9, R-scape 2.0.4, HMMER 3.4, BLAST+, Foldseek, MMseqs2, seqkit and ViennaRNA) (5, 25, 30, 56, 58, 86–88), and to the cluster scheduler.

### 4.34. Training and evaluation of embedding-based linear probes

We trained linear probes (40) on the centroids of L2-normalized Evo-PIN embeddings on each cluster (4,096 dimensions), with training labels derived from Rfam annotations. A cluster was considered annotated only when the same Rfam family overlapped its representative RNA motif in at least two distinct loci, excluding incidental matches elsewhere in the genomic region. The fluoride riboswitch probe used 4,808 annotated clusters across 29 families, with embeddings reduced to 256 principal components and evaluated against two negative sampling schemes: structured genomic backgrounds and flanking-region negatives (8,000 per fold), the latter controlling for genomic positioning characteristic of riboswitches. Both probes employed L2-regularized logistic regression and were evaluated using component and family hold-outs to assess generalization beyond closely related sequences and known RNA families. For the ribozyme probe, positive examples comprised 1,975 clusters spanning 16 Rfam families grouped into ten catalytic classes, including group II introns (841), RNase P (619), group I introns (275), hammerhead ribozymes (137), and six less abundant catalytic classes. To ensure label consistency, clusters with conflicting annotation assignments were excluded from the positive set, alongside group II intron fragments and ribosomal RNA fragments. Negative examples were drawn exclusively from known non-catalytic ncRNA families, including riboswitches, tRNAs, tmRNAs, SRP RNAs, 6S RNAs, and regulatory RNA elements. Of the 9,751 eligible clusters spanning 55 families, each family was subsampled to a maximum of 150 representatives, yielding 3,960 negatives and a combined training pool of 5,935 clusters. Unannotated clusters were never treated as negatives to avoid introducing potentially undiscovered catalytic RNAs into the background set. Sample weights balanced positive and negative classes and individual family contributions to prevent abundant families from dominating training.

The riboswitch probe achieved a median family hold-out AUROC of 0.997, with fluoride riboswitch classification remaining robust against both negative controls. Generalization to unseen ligand classes was limited (guanidine-I AP 0.087, near-random prevalence of 0.085). The hammerhead ribozyme probe achieved near-perfect component hold-out performance (AUROC 0.9995-0.9998), substantially outperforming a GC/length/taxonomy baseline (AUROC 0.50). Under family hold-out, recall at 1% FPR remained high for hammerhead (1.00), HDV-like (1.00), group I (0.996), and group II (0.94), but decreased for glmS (0.18) and RNase P (0.08). We also compared the performances of probes against cosine nearest-neighbor classifiers using identical embeddings and data splits. While performance was comparable under component hold-out, ribozyme probes substantially outperformed nearest neighbors on unseen families (hammerhead AUROC 0.9998 versus 0.672; other ribozyme nearest-neighbor AUROCs at ≤0.488). For fluoride riboswitches, the nearest-neighbor classifier’s performance remained closer to the probe (fluoride riboswitch AUROC 0.978 versus 0.997).

### 4.35. Rfam covariance model comparisons and sequence logos

Candidate clusters were evaluated against published Rfam CMs using Infernal 1.1.5 (58). The 307 sequences in the fluoride riboswitch candidate cluster (#418,142) were searched against the fluoride riboswitch CM on Rfam (RF01734) at maximum sensitivity. For ribozymes, 82 clusters (18,644 total sequences and 30 positive controls) were searched against 32 ribozyme and intron CMs in Rfam. Matches were required to exceed the model-specific Rfam gathering cutoff (GA), cover ≥80% of the model consensus length, overlap ≥50% of the candidate motif, and occur on the forward strand. No loci in the fluoride riboswitch candidate cluster #418,142 exceeded the RF01734 CM’s threshold within the noncoding motifs of the cluster (maximum 28.9 bits; GA 34.0). Similarly, for the hammerhead candidate cluster #1,533,398, none of 152 loci passed the thresholds of any of the six hammerhead CMs (RF00163, RF02276, RF00008, RF02275, RF02277, RF03152) with a best hit having the score of 26.4 bits against RF00163’s GA of 34.0. To compare the novel divergent motifs with the canonical motifs, the sequences mapped to the novel motif and the Rfam seed sequences were independently aligned to the same covariance model for the fluoride riboswitch family (RF01734) and a hammerhead family (RF00163) using cmalign. Alignments were restricted to consensus columns, and unweighted sequence logos were generated using the information content of each column with small-sample correction. The fluoride riboswitch comparison included 306 candidate loci and 289 Rfam seed sequences across 64 consensus positions; the hammerhead comparison included 142 candidate loci and 29 seed sequences across 45 positions.

### 4.36. In silico saturation mutagenesis of divergent hammerhead and fluoride riboswitch motifs

To identify nucleotide positions contributing to probe predictions, we performed *in silico* saturation mutagenesis (ISM) (41, 42). For each case, we tested a single representative locus, selected as the sequence with the length that is closest to the median representative locus length. This yielded one locus from 307 representative loci for the fluoride riboswitch candidate (cluster #418,142) and one from 152 for the hammerhead-like candidate (cluster #1,533,398). We extracted Evo-PIN embeddings from each locus, applying substitutions only at the unpadded positions (not the flanking padding positions), giving 228 positions and 684 variants for the fluoride riboswitch locus and 85 positions and 255 variants for the hammerhead locus, with every variant yielding a finite score from the respective linear probe. Sensitivity of the probe to base changes was quantified as the difference between the wild type and mutant probe score. The per-position score was calculated as the mean of the probe score differences resulting from the three possible substitutions. Positions were then highlighted using annotation established independently of the probe – the RF01734-aligned region for the fluoride riboswitch locus and the conserved positions of the CUGANGA and GAAA motifs for the hammerhead locus.

### 4.37. Evidence rubric for investigating nuclease/helicase domain containing proteins

Candidate prioritization followed a predefined rubric targeting clusters of unannotated structured RNAs colocalized with proteins containing both nuclease and helicase domains. Nine weighted criteria (scored 0-4) assessed the RNA architecture, colocalized protein identity, genomic association, sequence conservation, taxonomic breadth, covariation within the RNA structure, novelty, and other distinctive features. Missing evidence contributed zero to the lower-bound score, with evidence coverage tracked separately. Candidate retention required a score ≥60%, evidence coverage ≥80%, and passage of eleven mandatory checks. The campaign comprised four independent agent rollouts, each screening candidates using Pfam domain annotations and Foldseek structural context, with separate provenance records and no prior family hints. Independent reviewer agents, blinded to the discovery process, re-evaluated protein-domain assignments, genomic context, Rfam covariance model hit overlaps, alternative annotations, and supporting literature. Final rankings were computed deterministically from reviewed evidence.

### 4.38. Analysis of publicly available RNA-seq data

Nine publicly available paired-end RNA-seq libraries from *Pedobacter mucosus* Q8-18 (BioProject PR-JNA1244275; SRR32921415–SRR32921423) were retrieved from SRA, quality-trimmed using fastp (89), and aligned to the reference chromosome (GCF_022200785.1) using Bowtie2 (90). Of 121.7 million read pairs, 120.9 million passed quality filtering, with alignment rates of 98.56-99.34%. High-confidence alignments were used to calculate per-base fragment coverage, counting overlapping paired mates once. Library strandedness was inferred from annotated coding sequences, with all nine libraries exhibiting R1-antisense orientation. Expression was quantified as the within-library ratio of mean sense-strand coverage across the 357-nt candidate RNA motif to the adjacent *phoH2* coding region, with coverage tracks normalized per million cleaned read pairs.

### 4.39. Phylogenetic investigation of PhoH2 proteins

PhoH2 proteins carrying both PIN_4 (PF13638) and PhoH (PF02562) domains were identified by using existing protein annotations built in previous steps, where Pfam architectures were assigned to all 1.47B proteins neighboring the intergenic regions in our dataset. Selection required both domains to be called on the same protein identifier and was not restricted to any cluster subset, yielding 88,404 PhoH2 proteins flanked by 46,714 intergenic loci immediately on the 5^′^ side and 48,286 on the 3^′^ side. For each intergenic locus on the 5^′^ side, the flanking PhoH2 gene was mapped to its MMseqs representative and the representative sequence retrieved, with genomic genes lacking a representative recovered from their assembly’s protein FASTA; 31,025 of 44,715 loci (69%) resolved to a protein, collapsing onto 7,477 distinct representative sequences. The remaining 31% are metagenomic genes. Sequences were aligned using MAFFT v7.525 (69), trimmed with trimAl to 413 informative columns (91), and used to infer a maximum-likelihood phylogeny with FastTree 2.2.0 under the LG + Γ model (92). The tree was displayed unrooted. Independently, proteins were grouped using MMseqs2 at 50% sequence identity and 80% coverage, yielding 792 groups, of which 51 contained at least 30 loci. Group membership was mapped onto the phylogeny for visualization. Associated RNA families were identified independently of protein sequences, enabling assessment of RNA–protein evolutionary associations without circular classification.

## 5. Acknowledgements

We thank all members of the Hsu and Konermann labs for advice and feedback. We thank C. Herrmann for helpful discussions, J. Sullivan and S. Newins for managing the Arc computing infrastructure used for this project, J. Tey and I. Mukherjee for advice on categorical jacobians and fine-tuning methods, and J. Jiao and S. Eddula at OpenAI for advice on agentic research workflows.

## Author contributions

G.S., T.H., M.G.D., and P.D.H. conceived the study. A.S.P., S.K., and P.D.H. supervised the research. G.S., T.H., J.C.G., G.A.G., Z.M., and F.W. developed the workflow and performed computational analyses. C.R.-T., G.S., T.H., and J.C.G. designed the visualizations. P.Y., T.H., and G.A.G. built the accompanying software. G.S., T.H., J.C.G., A.P., A.S.P., and P.D.H. wrote the manuscript with input from all authors.

## Data, code, and materials availability

COMPASS outputs from this work, including the assignment of sequences into the 5.14M clusters, CMfinder motif alignments of each cluster, colocalized gene annotations, and Evo-PIN token entropy tracks, are available through the Model Context Protocol (MCP) server (https://compass.arcinstitute.org/mcp). Data for a subset of 255,213 clusters, selected based on strong structural support for their noncoding motifs and the information in their genomic context (see Methods), are also available through an interactive web app (https://compass.arcinstitute.org/).

## 6. Supplementary Figures

**Fig. S1.**
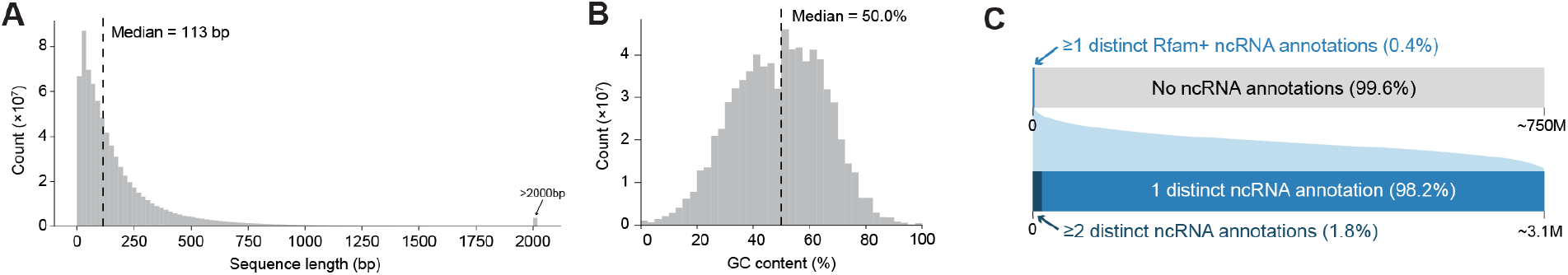
Properties of 750M representative intergenic regions parsed from prokaryotic genomes and metagenomes. (A) Distribution of the lengths of the intergenic sequences. (B) Distribution of the GC content of the intergenic sequences. (C) Fractions of sequences with significant hits to covariance models of known ncRNAs (top), and the fraction of sequences with hits that have hits from ≥2 distinct ncRNAs (bottom).

**Fig. S2.**
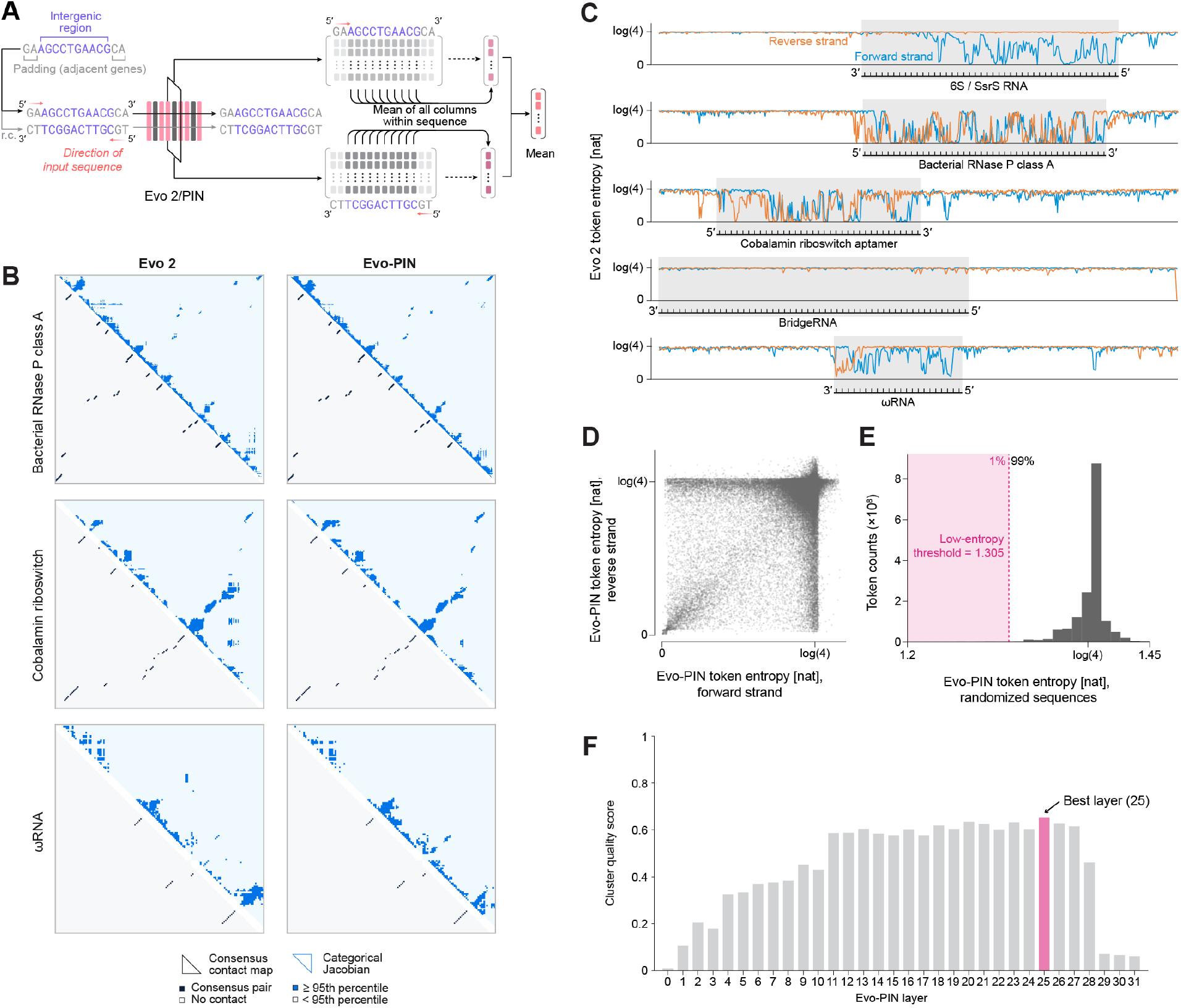
Methods for extracting pooled embeddings were tested and optimized on a subset of intergenic sequences. (A) Schematic diagram of the simple mean pooling function. (B) Categorical Jacobians of Evo 2 and Evo-PIN for additional ncRNA families. Family-averaged contact maps for bacterial RNase P class A, the cobalamin riboswitch and *ω*RNA, binarized at the 95th percentile (upper triangle) and compared with the consensus base pairs of each family’s covariance model (lower triangle), as in Fig. 1D. Consensus columns without aligned test sequences are omitted. (C) Token entropies of Evo 2 on intergenic sequences with known ncRNAs showed that positions along the sequences that were part of known ncRNAs had lower token entropies, but less consistently compared to Evo-PIN. (D) Comparison of Evo-PIN token entropies for the same position of intergenic sequences calculated using context from the forward strand versus the reverse strand. For many positions, these two token entropies are distinct. (E) Distribution of Evo-PIN token entropies for randomized sequences of matching lengths to the intergenic regions. The lower 1% percentile was chosen as the threshold for low-entropy pooling of embeddings. (F) Comparison of cluster quality scores for low-entropy pooled embeddings extracted from each block of Evo-PIN.

**Fig. S3.**
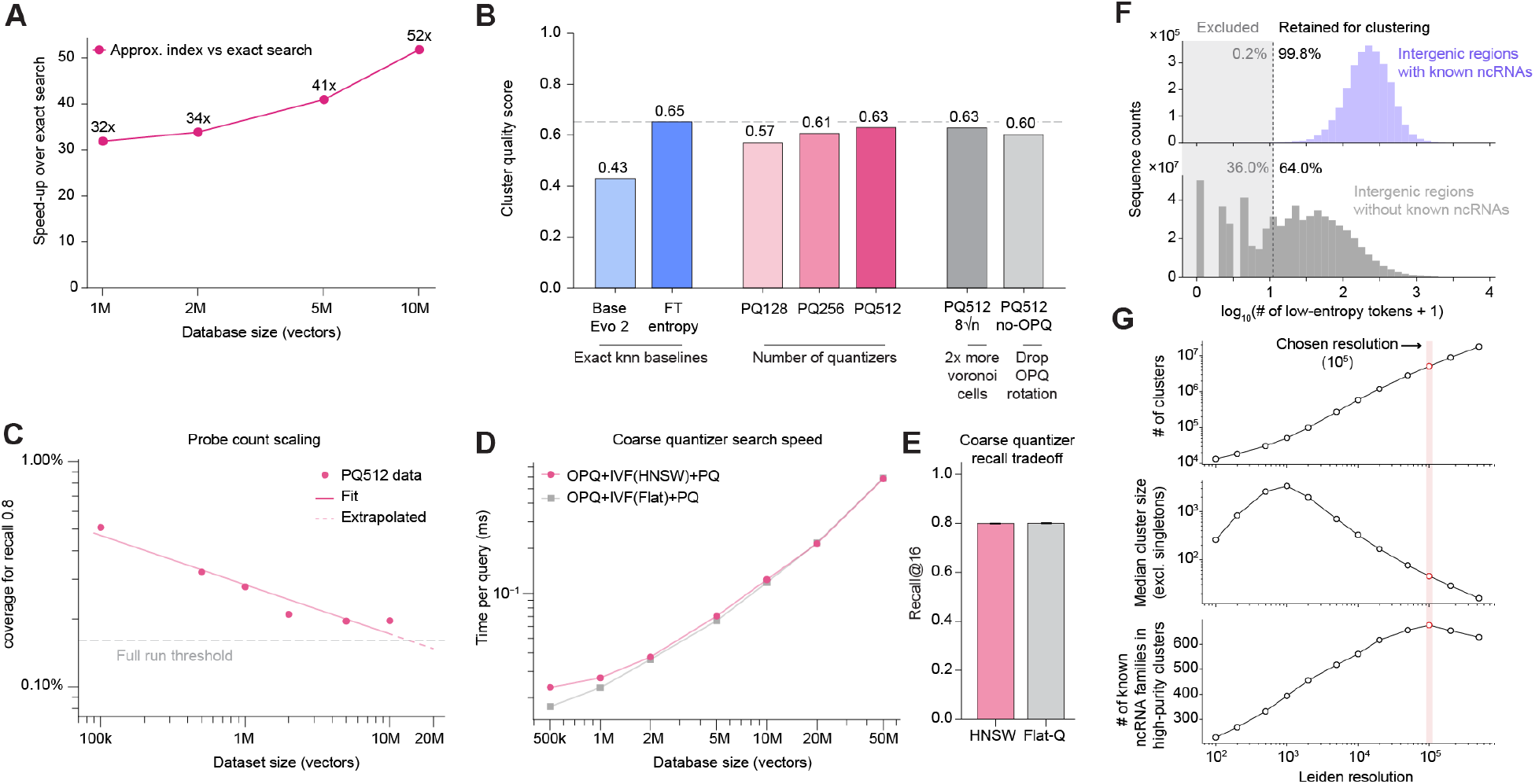
Experiments to optimize the approximate nearest neighbor search and the Leiden clustering used to cluster the pooled embeddings at scale. (A) Relative speed-up of final approximate search method compared to exhaustive search, measured for different database sizes. (B) Comparison of cluster quality scores for different approximate kNN calculation strategies to scores of exact kNN baselines. (C) Percentage of IVF cells that needed to be searched to reach 0.8 recall compared to exact search. The gray dashed line shows the coverage that was chosen for the full clustering of pooled embeddings. (D) Comparison of the scaling of search time per query over IVF centroids with exact and approximate methods (coarse quantizer). (E) Neighbor recall of coarse quantization methods compared to exact search for a 10M vector database. (F) Distributions of the number of low-entropy tokens for sequences with known ncRNAs compared to sequences without known ncRNAs. Sequences with less than 10 low-entropy tokens were excluded from clustering, which removed 36.0% of sequences without ncRNAs, but only 0.2% of sequences with ncRNAs. (G) Across multiple Leiden resolutions, we compared the cluster counts, median cluster sizes, and the number of known ncRNA families that are grouped into high-purity clusters with ≥50 sequences. The resolution of 10^5^ was chosen to optimize for the number of known ncRNA families in high-purity clusters.

**Fig. S4.**
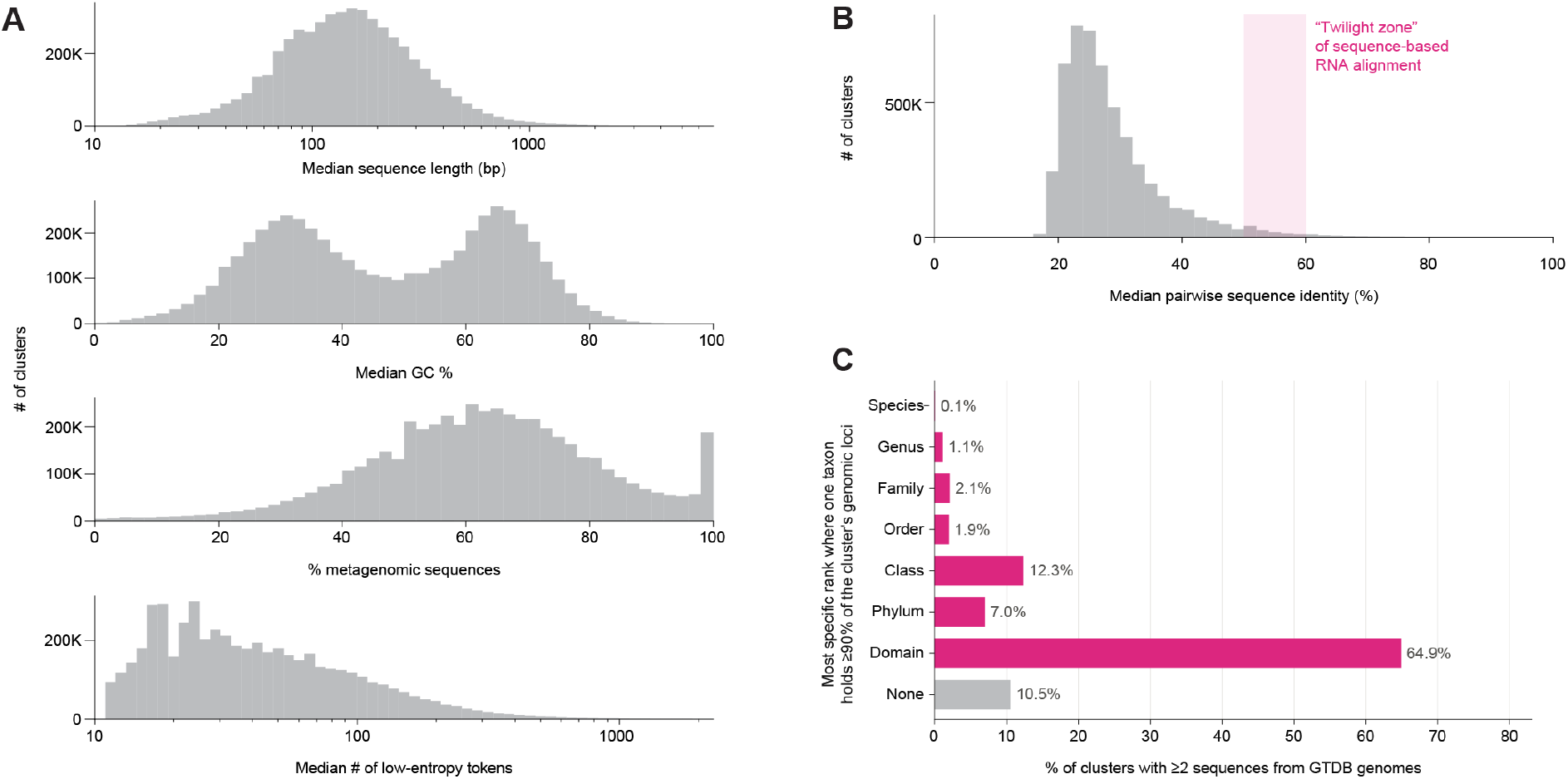
General properties of 5.14M clusters of intergenic regions. (A) Distributions of the median sequence length, median GC%, % of metagenome-derived sequences, and the median number of low-entropy tokens across clusters. (B) Distribution of the median pairwise sequence identities within each cluster. For most clusters, this value falls below the “twilight zone” of sequence-based RNA alignment where the alignment is reliable. (C) Taxonomic distributions of the sources of intergenic sequences parsed from genomic sequences in each cluster. Every sequence prior to deduplication at 70% sequence identity was counted for each cluster.

**Fig. S5.**
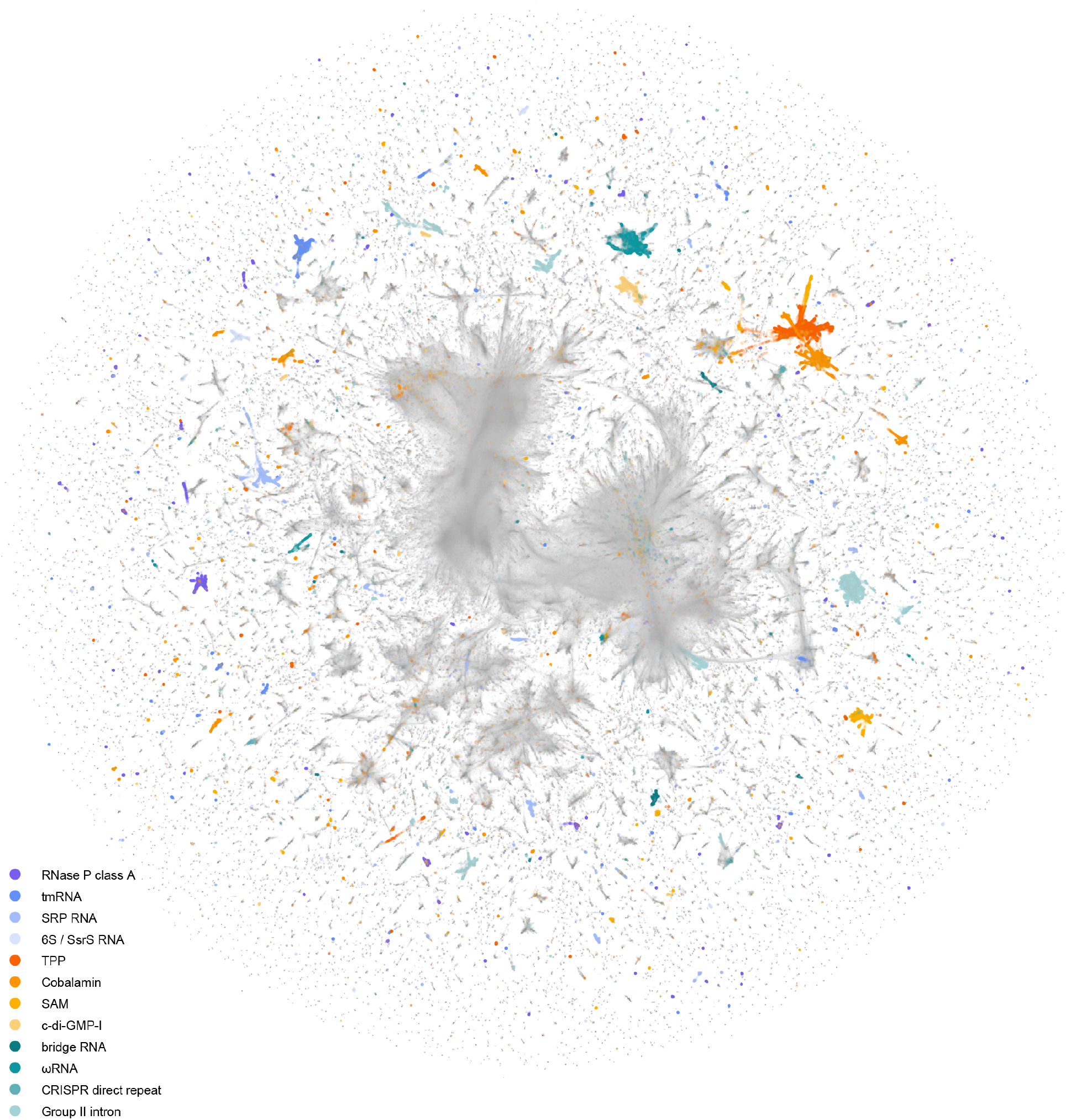
UMAP visualization of COMPASS clusters with known noncoding RNAs. An expanded version of the UMAP visualization in Fig. 1B. Sequences containing 12 known representative ncRNAs are highlighted.

**Fig. S6.**
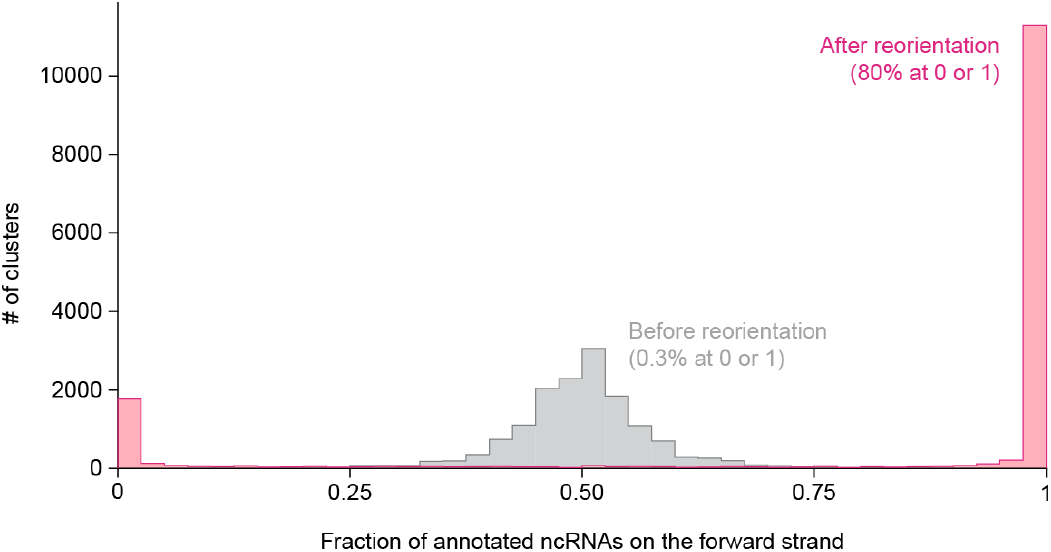
Testing the embedding-based strand reorientation method on COMPASS clusters of known ncRNAs. Distribution of the fraction of annotated ncRNAs that are on the forward strand after applying the embedding-based strand reorientation method on clusters of known ncRNAs. For 80% of the clusters, this method resulted in a perfectly consistent orientation, where all annotated ncRNAs in the cluster were situated on the same strand.

**Fig. S7.**
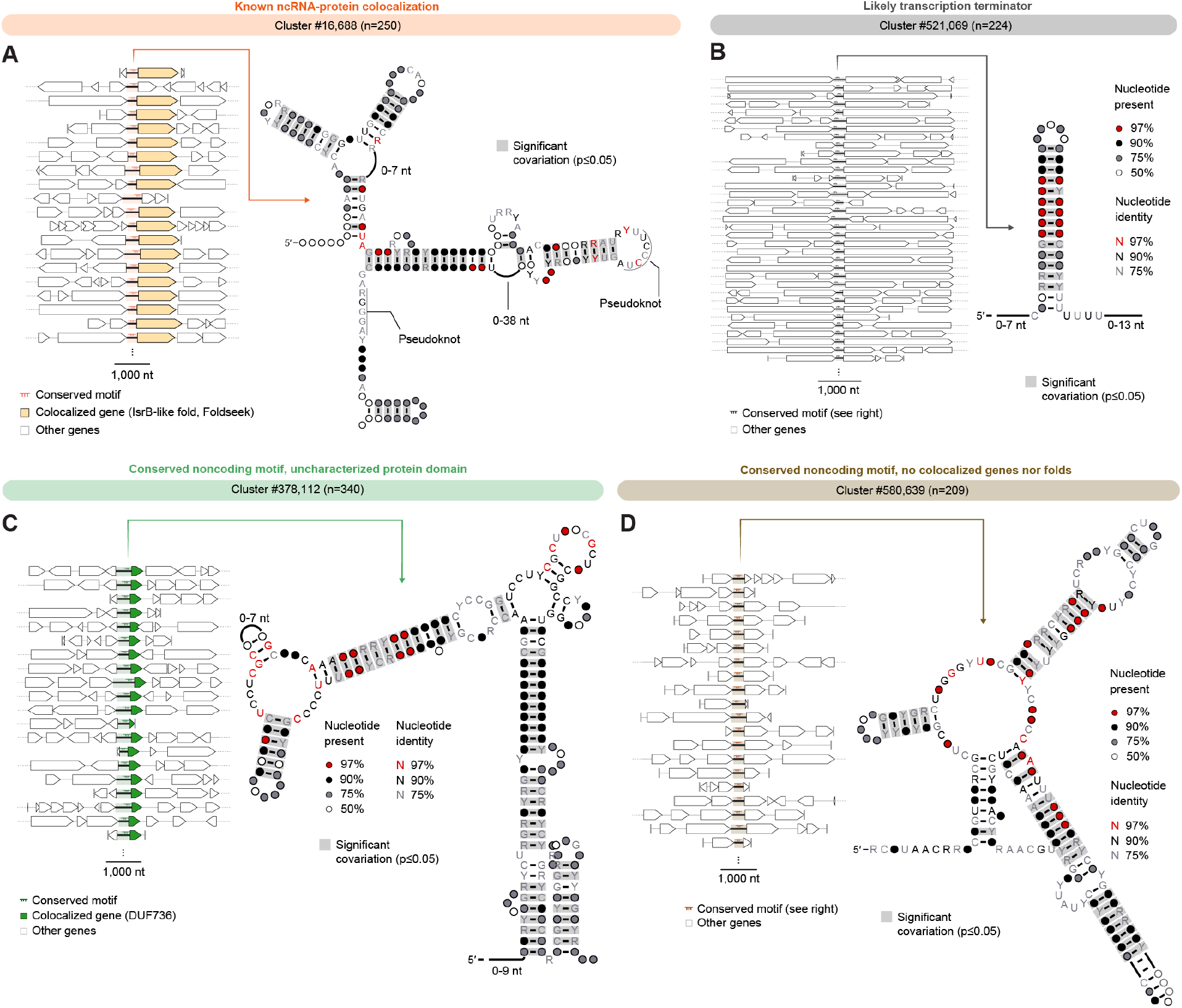
Additional examples of COMPASS clusters with shared noncoding motifs, and their genomic contexts. (A) Sequences of cluster #16,688 share a motif upstream of a colocalized, IsrB-like fold (left). The motif displays known structural hallmarks of *ω*RNAs including the nexus pseudoknot (right). (B) Sequences of cluster #521,069 share a motif that lies downstream of flanking protein-coding genes, without a significantly colocalized protein family in its context (left). The motif’s structure resembles the known structure of rho-independent transcriptional terminators, with a hairpin stem followed by a string of U’s (right). This cluster/motif was classified as a likely transcription terminator. (C) Sequences of cluster #378,112 share a motif upstream of a colocalized gene that encodes for an uncharacterized protein (DUF736). (D) Sequences of cluster #580,639 share a motif, but do not significantly colocalize with any gene families nor folds.

**Fig. S8.**
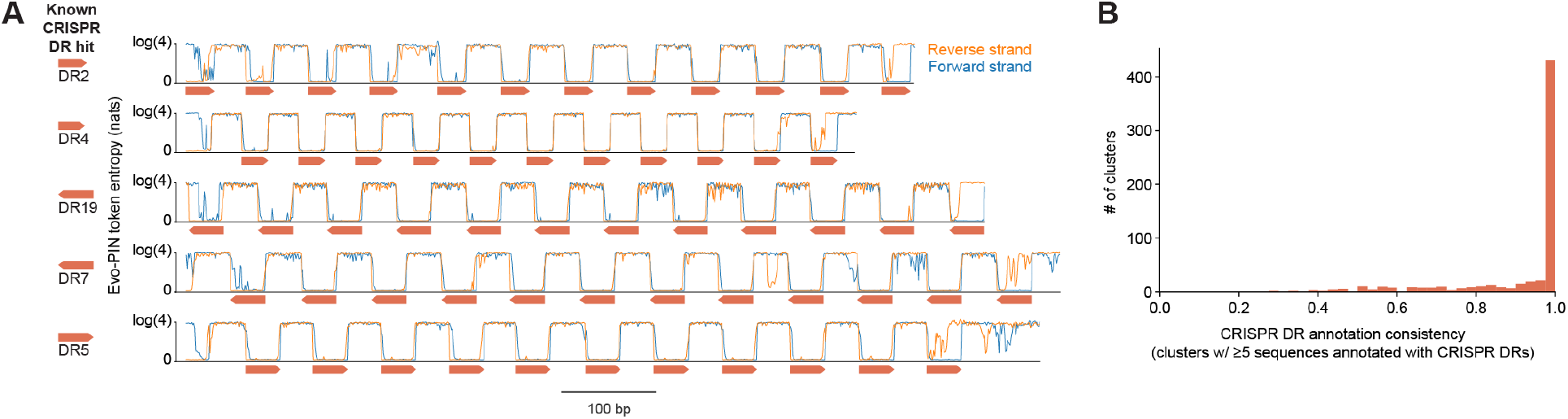
Known CRISPR arrays display distinct Evo-PIN token entropy patterns and cluster together with pooled embeddings. (A) Evo-PIN’s token entropies on sequences with CRISPR arrays, alongside the annotations of the sequences with known CRISPR direct repeat models. The entropies display a periodic pattern that coincides with the structure of the array. (B) Distribution of the consistencies (fraction of the sequences annotated with the most common annotation) of CRISPR direct repeat annotations among the 621 clusters with ≥5 sequences uniquely annotated with a known CRISPR direct repeat.

**Fig. S9.**
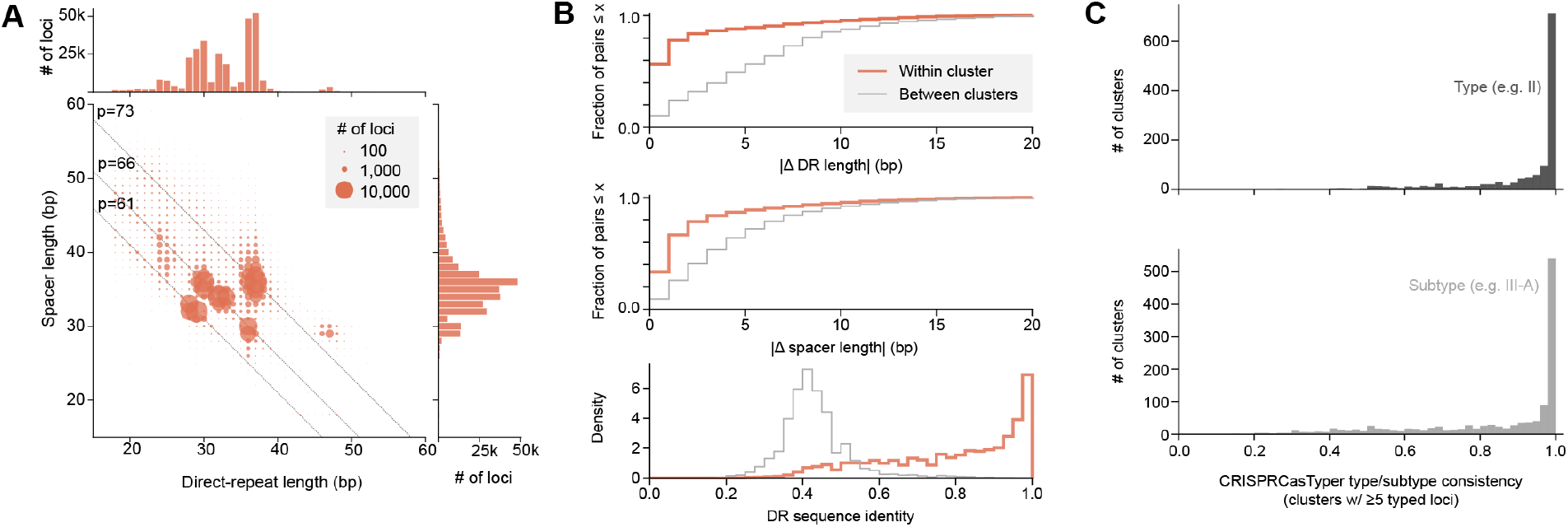
Properties of CRISPR array-like clusters identified from COMPASS. (A) Distributions of the DR and spacer lengths of individual loci within the CRISPR array-like clusters. (B) Comparison of the lengths of the direct repeat (DR) and spacer, and the sequences of the DR, between sequences within the same cluster and in different clusters. Sequences within the same cluster had more consistent DR/spacer lengths and DR sequences. (C) Distribution of the consistencies (fraction of the sequences annotated with the most common type/subtype) of CRISPR system type/subtype annotations among the CRISPR array-like clusters with ≥5 typed loci.

**Fig. S10.**
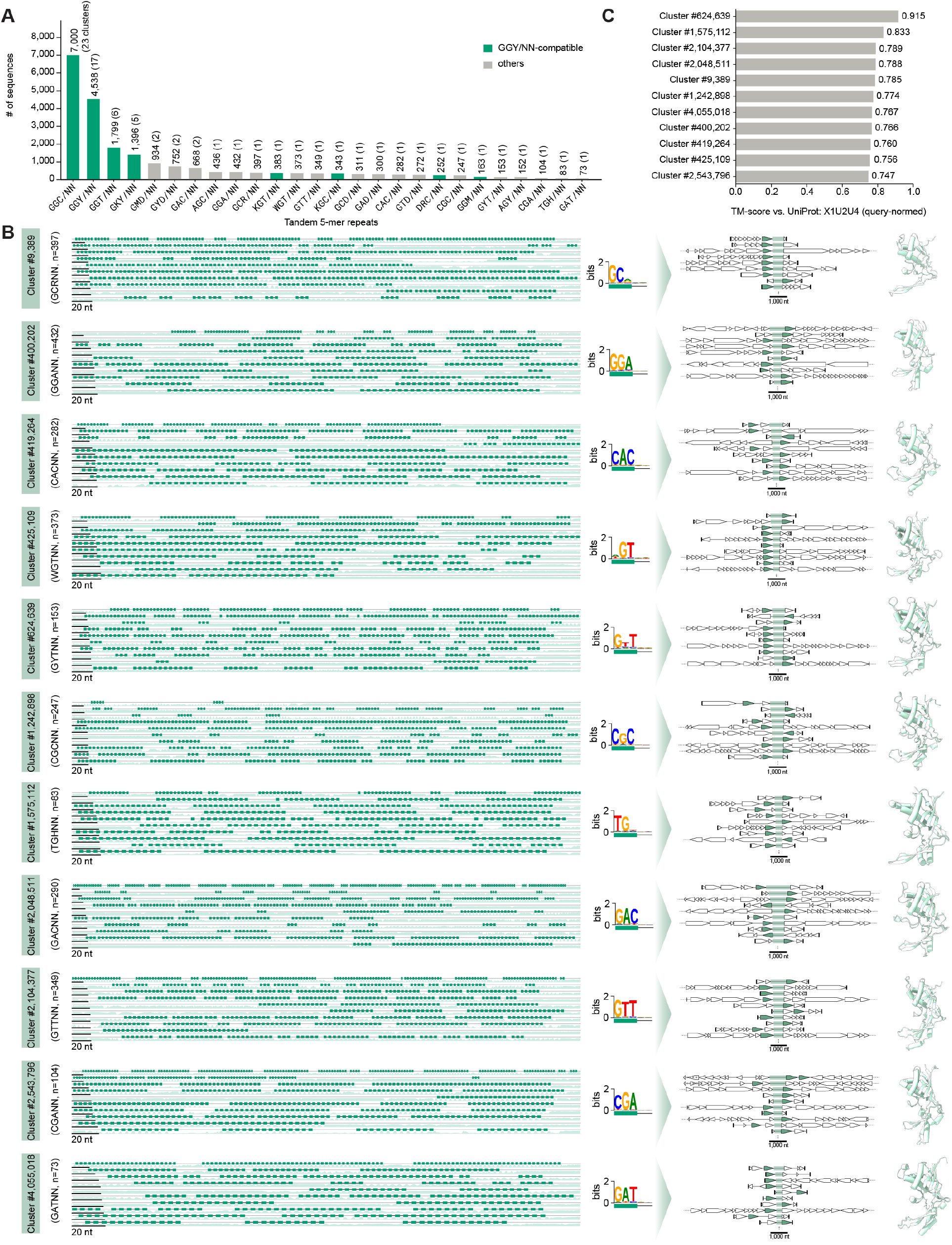
VIPR arrays with noncanonical repeat patterns. (A) Total number of sequences in each tandem 5-mer repeat pattern observed in VIPR array-like clusters. Bars for patterns that are compatible with the canonical GGY/NN pattern are highlighted in green. (caption continued on next page) (B) The repeat array structures, the sequence logo of the 5-mer repeat, and the genomic context of 11 VIPR array-like clusters with noncanonical repeat patterns. The conserved protein folds resembling the reported Vipr protein (UniProt: X1U2U4) are highlighted in the genomic context. ESMFold-predicted structures of the representative proteins of the conserved fold are shown on the right. (C) Query-normed TM-scores of the conserved Vipr-like folds colocalized in the 11 clusters against the reported Vipr protein (UniProt: X1U2U4).

**Fig. S11.**
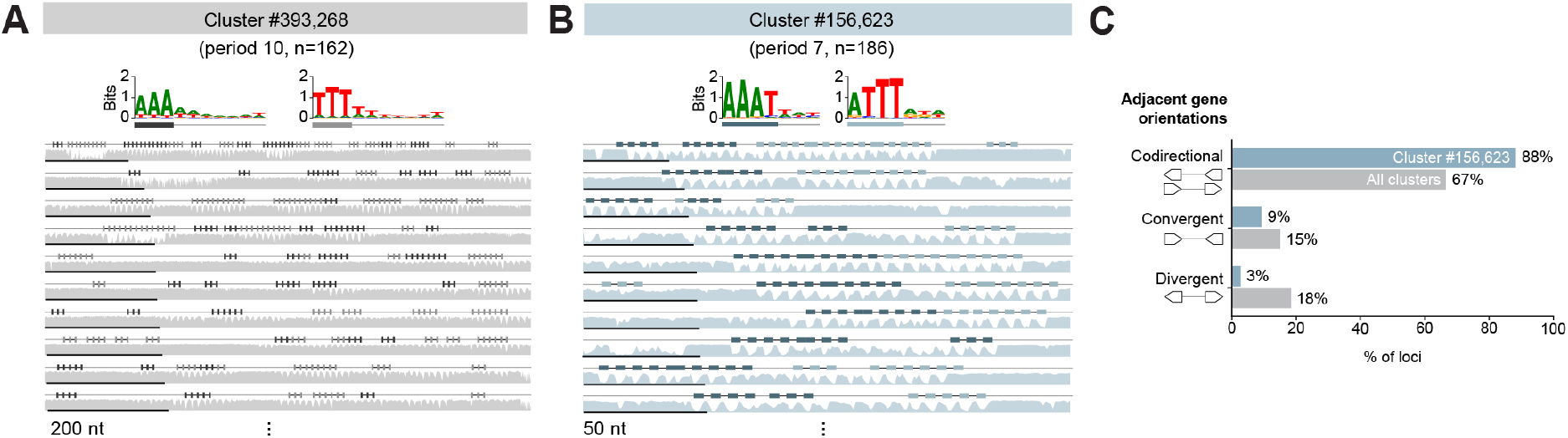
Properties of clusters with token entropy periods that fall outside the ranges of CRISPR, TIGR, and VIPR arrays. (A) Cluster #393,268 (n=162), with a dominant period of 10 nt, displayed as in Fig. 3C. The loci in this cluster contained runs of 4-6 adenines or thymines spaced every 10-11 nt, consistent with A-tracts associated with intrinsic DNA curvature rather than structured RNA elements. (B) Cluster #156,623 (n=186), with a dominant period of 7 nt, displayed as in Fig. 3C. The two sequence logos show the tetramers of the two arrays. (C) Comparison of the relative orientation of the two genes flanking each locus for cluster #156,623 with all clusters.

**Fig. S12.**
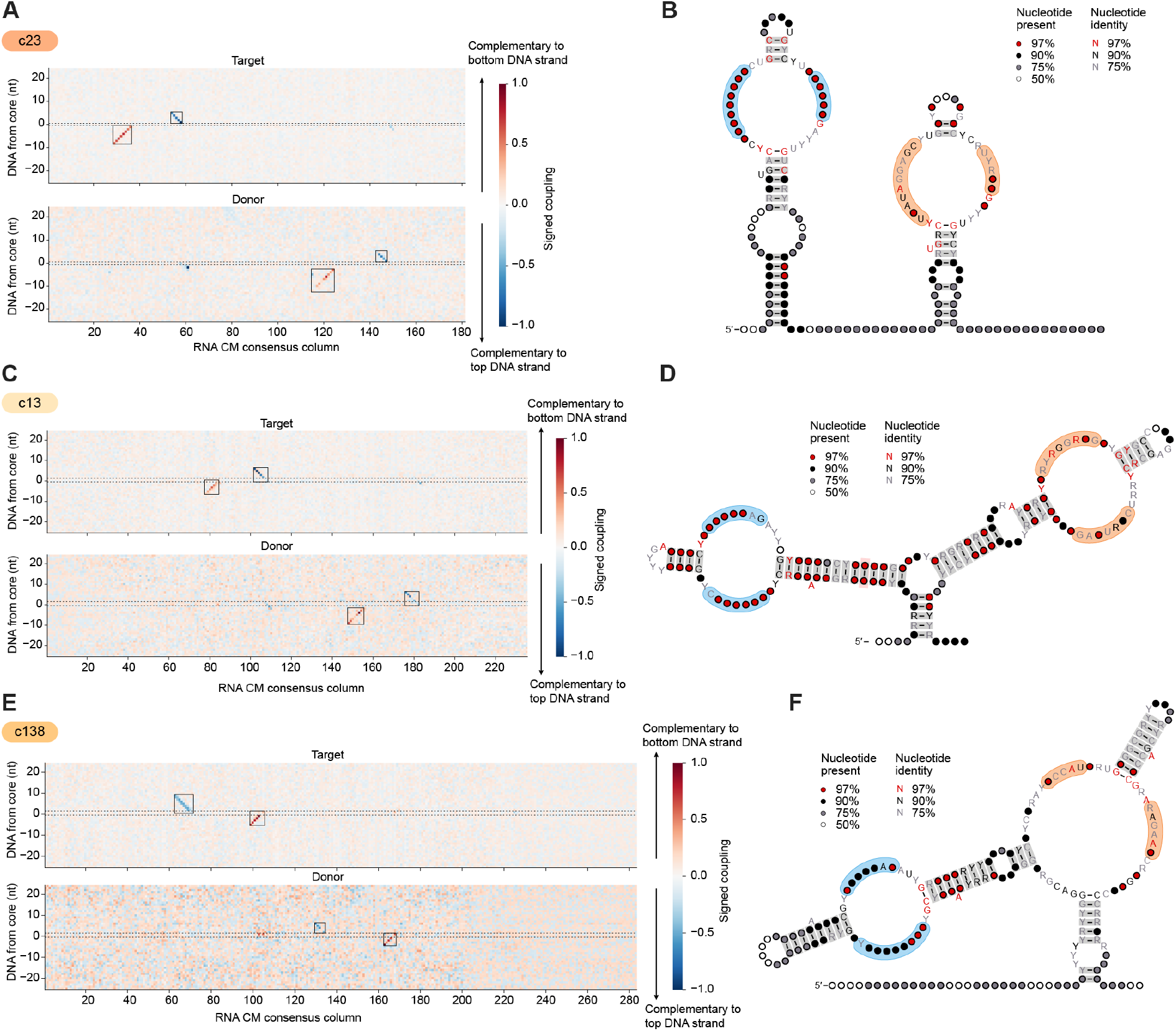
Bridge RNAs with diverse secondary structures show guide loop covariation with their target and donor DNA. (A) Covariation between bRNA and DNA positions for cluster 23. Top: signed coupling between each RNA consensus column and each DNA position relative to the core, for the target site. Bottom: signed coupling for the donor site; positive and negative values indicate complementarity to the bottom and top DNA strand, respectively. Boxes mark regions with strong covariation between bRNA and DNA. (B) Consensus secondary structure of the cluster 23 bRNA, with regions with strong covariation shaded. Circle fill indicates the fraction of sequences with a nucleotide at that position, and letter color indicates nucleotide conservation. (C) As in (A) for cluster 13. (D) As in (B) for cluster 13. (E) As in (A) for cluster 138. (F) As in (B) for cluster 138.

**Fig. S13.**
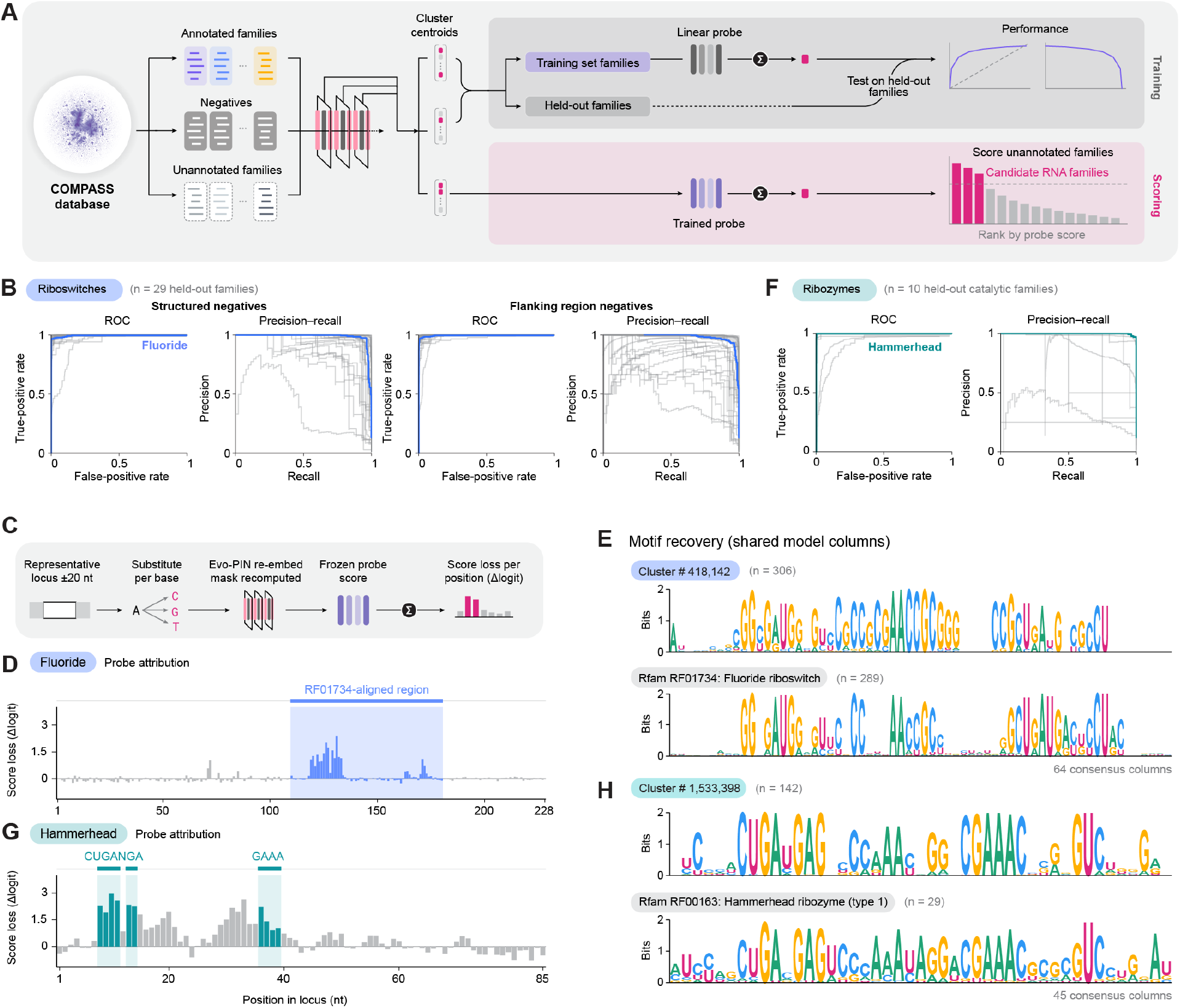
Training, evaluation, and sequence-level interpretation of Evo-PIN embedding-based ncRNA probes. (A) Schematic of linear probe training on Evo-PIN cluster embeddings, with held-out family evaluation followed by scoring of unannotated clusters. (B) ROC and precision–recall curves for 29 held-out riboswitch families against structured and flanking-region negatives. Fluoride riboswitch performance is highlighted in blue. (C) Position-wise *in silico* mutagenesis measuring changes in probe logits following nucleotide substitutions and re-embedding. (D) Attribution profile highlighting the canonical fluoride riboswitch region. (E) Sequence logos comparing the consensus residues of the candidate cluster #418,142 and the existing Rfam fluoride riboswitch family (RF01734). (F) ROC and precision–recall curves for ten held-out catalytic RNA families, with hammerhead ribozyme performance highlighted in teal. (G) Attribution profile highlighting conserved hammerhead catalytic motifs, as in (D). (H) Sequence logos comparing candidate #1,533,398 with respective Rfam hammerhead (RF00163) family. Sequences were aligned to shared covariance model consensus positions to compare nucleotide conservation.

**Fig. S14.**
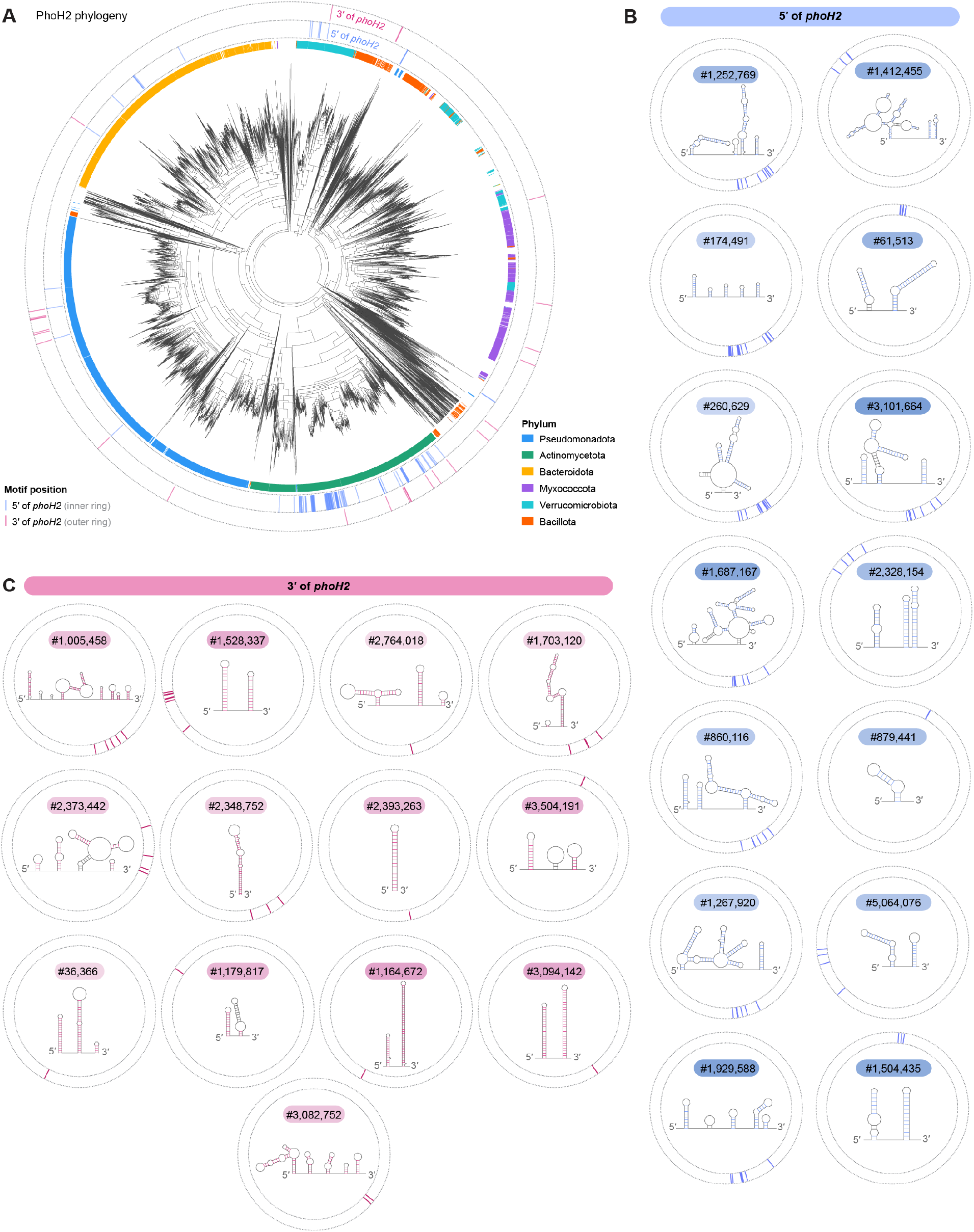
Phylogenetic diversity of PhoH2-associated structured ncRNAs. (A) Unrooted maximum-likelihood phylogeny of 7,477 representative PhoH2 (PIN_4–PhoH) proteins, annotated by bacterial phylum (inner colored ring) and associated clusters with noncoding motifs located 5^′^ (blue) or 3^′^ (pink) of *phoH2* (outer tracks). RNA clusters were identified independently of protein phylogeny and subsequently mapped onto the tree, revealing lineage-specific associations between PhoH2 homologs and distinct structured RNAs. (B) Predicted consensus secondary structures of the noncoding motifs of 14 PhoH2-associated clusters located on the 5^′^ side of *phoH2*. Each structure is labeled by the cluster identifier, with blue marks on the surrounding circular track indicating the phylogenetic distribution of associated PhoH2 proteins in (A). (C) Predicted consensus secondary structures of the noncoding motifs of 13 clusters located on the 3^′^ side of *phoH2*. Pink marks on the surrounding circular tracks indicate their corresponding phylogenetic distributions in (A).

